# Beyond the 1:1 Ligand–Protein Paradigm: An In Silico Assay for Competitive Ligand Binding

**DOI:** 10.1101/2025.10.31.685849

**Authors:** Vince Bart Cardenas, Stefano Raniolo, Paolo Conflitti, Vittorio Limongelli

## Abstract

Competitive binding assays (CBAs) are widely used in drug discovery to quantify and compare ligand/receptor affinities. However, their molecular interpretation is often limited by the inherent complexity of ligand–receptor interactions and the transient nature of binding events. All-atom molecular dynamics (MD) simulations offer valuable mechanistic insights in ligand binding studies, but remain computationally prohibitive for capturing the long-timescale, multiligand behavior characteristic of CBAs. Therefore, characterizing the molecular aspects of CBAs remains a fundamental challenge in molecular biophysics. Here, we introduce a coarse-grained MD (CGMD) approach capable of recapitulating CBA-like dynamics between ligands of opposing efficacy— the full agonist NECA and the inverse agonist ZM241385— at the adenosine A_2A_ receptor, a prototypical G protein-coupled receptor and a key pharmacological target. By simulating ligand mixtures at varying molar ratios, we capture hallmark features of experimental CBAs, including spontaneous binding, unbinding, and direct competition at the orthosteric site. Our simulations reveal an extracellular vestibular site that modulates ligand access to the binding pocket. Occupation of this site by NECA facilitates ZM241385 entry and prolongs its residence time, revealing a cooperative mechanism within an otherwise competitive process. These findings offer a molecular perspective on ligand competition at GPCRs and demonstrate the potential of CGMD as a viable method for probing multiligand dynamics in drug discovery.

## 1 Introduction

Competitive binding assays (CBAs) are a cornerstone of drug discovery and development, providing fundamental quantitative metrics that characterize the interaction between ligands and their molecular targets.[1–4] In high-throughput screening campaigns, for example, CBAs are routinely employed in the identification of promising lead compounds. [5, 6] In a typical CBA, the binding of a labeled ligand to a target molecule is measured at equilibrium. A second, unlabeled ligand is then introduced to compete for the same binding site, displacing the labeled ligand in proportion to its own affinity. By quantifying this displacement, critical properties such as binding affinity and inhibitory potency can be derived. [7–9]

Since inception in 1959, CBAs have diversified and improved in scope and methodology. [10, 11] However, several limitations remain — from technical challenges like radioactive tracer use to more serious concerns like assay conditions inadvertently altering ligand pharmacological profiles. [12, 13] Furthermore, interpreting CBA results to gain meaningful insights into ligand molecular traits is anything but straightforward. [14] At their core, CBAs measure macroscopic observables — quantities that emerge from complex molecular processes that are often hidden from view. These include protein conformational changes as well as transient and more stable ligand–protein and ligand–ligand binding interactions governed by non-covalent forces, the mechanistic details of which frequently remain unknown. [7, 15] The influence of such molecular-level dynamics on macroscopic assay outcomes is undeniable, yet their direct experimental characterization is extremely difficult, if not entirely impossible. As a result, our current understanding of competitive binding is incomplete at best or severely limited at worst — particularly when viewed through a molecular lens. Bridging this gap requires tools that can resolve molecular phenomena at atomic resolution and capture the rapid dynamics inherent to binding processes. Fortunately, this challenge can be embraced by the computational sciences, particularly through the use of classical molecular dynamics (MD) simulations.

For decades, MD simulations have allowed scientists to observe molecular behavior in near-atomic detail and in a time-resolved fashion, capturing the fundamental forces that govern molecular interactions. These simulations have revealed mechanistic, structural, thermodynamic, and kinetic insights into a wide range of protein-ligand systems, significantly advancing both molecular biology and drug discovery. [16–22] However, multiligand MD studies remain rare. Even among the few available, ligands are typically already initialized in or near the binding site, effectively bypassing the true complexity of ligand exploration and competition. [23–26] To authentically replicate a CBA environment in silico, simulations must allow ligands to diffuse freely, explore the protein surface, interact dynamically among themselves, and compete — or even cooperate — for binding sites. This demands long timescales, significant conformational sampling, and large simulation ensembles. With conventional all-atom molecular dynamics (AAMD), such simulations are simply computationally unfeasible due to the required time and resource investment. As a result, an atomistic simulation of a CBA is yet to be realized.

In this work, we present a proof-of-concept study that successfully emulates CBA-like conditions using a fully unbiased, multiligand coarse-grained molecular dynamics (CGMD) protocol. By foregoing AAMD for CGMD, we overcome critical timescale limitations thanks to CGMD’s computational efficiency and accelerated sampling, which arises from its simplified representation of the molecular system. In CGMD, groups of atoms are modeled as single interaction sites or “beads”, resulting in an overall fewer particle count, fewer computations per time step, and a smoother potential energy surface (PES).[27–29] Our target system is the adenosine A_2A_ receptor, a prototypical member of the largest and pharmacologically relevant protein family in eukaryotes G protein-coupled receptors (GPCRs) — encoded by approximately ~4% of the human genome [30, 31] and targeted by over one-third of all marketed drugs. [16, 32, 33] In particular, *we conducted five distinct in silico CBAs, each featuring a different molar ratio of two ligands: the full agonist NECA (Fig. 1a) and the inverse agonist ZM241385 (ZMA, Fig. 1b)*. By modulating ligand ratios, we mimic the effect of varying concentrations in experimental CBAs. Collectively, our unbiased CGMD simulations reach the millisecond timescale — equivalent to second-to-minute real time considering the CGMD sampling acceleration.

**Fig. 1.**
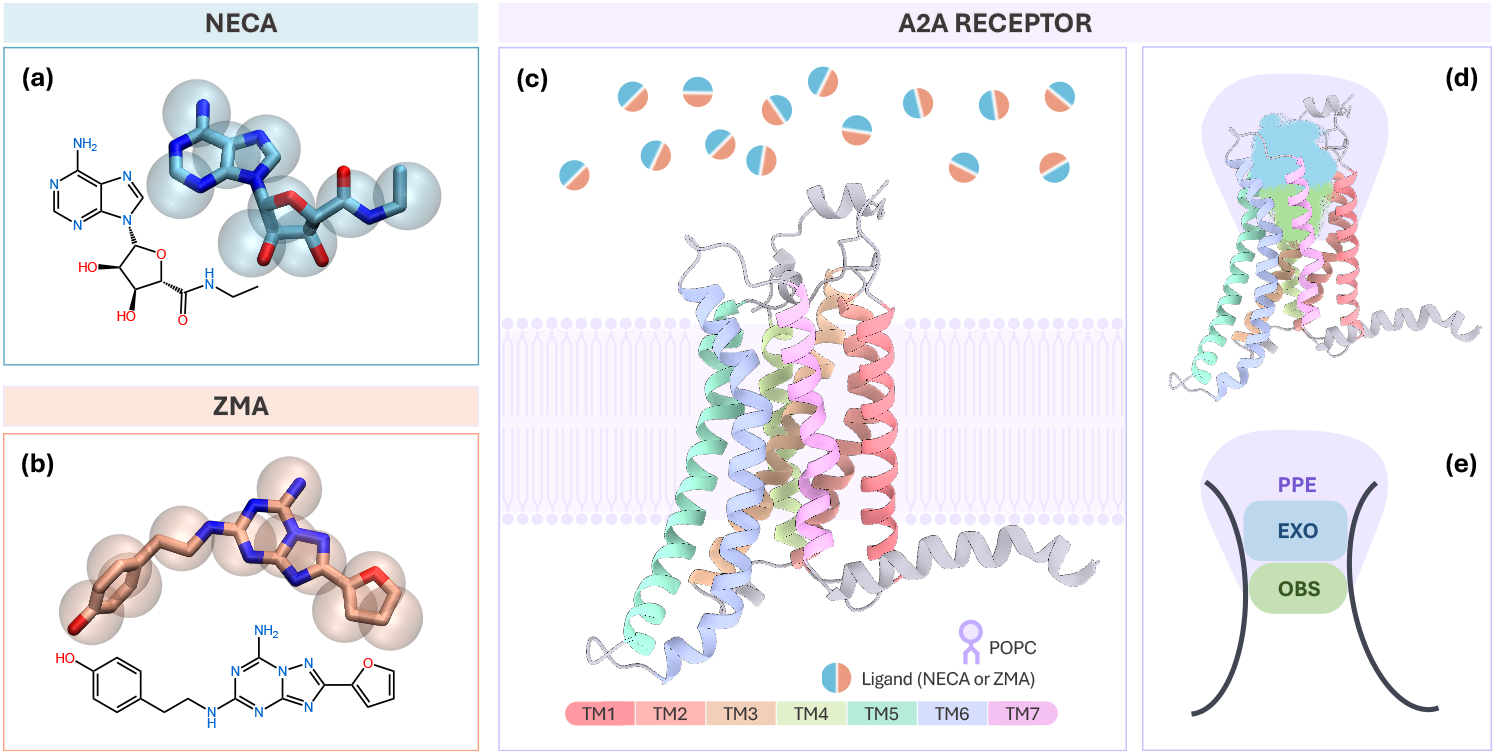
Orthosteric ligands NECA and ZMA and the A_2A_ adenosine receptor. **(a-b)** Chemical structures of the agonist NECA (blue) and the inverse agonist ZMA (orange), overlaid with their coarse-grained bead representation. **(c)** Ribbon representation of the A_2A_ receptor, showing the seven transmembrane helices (rainbow colors) and the ensemble of extracellular ligands (dual-color balls). **(d)** Point-cloud maps of ligand positional data, identifying the orthosteric site (light green), the exosite (light blue), and the immediate extracellular region surrounding the receptor (light purple). **(e)** Schematic overview of the receptor and its three spatial regions, which define the reference compartments used throughout the study.

Throughout the simulations, we observed multiple spontaneous binding and unbinding events of both NECA and ZMA to A_2A_, allowing us to extract mechanistic, kinetic, and thermodynamic insights. Notably, we observed a structural feature of A_2A_ represented by an antechamber region preceding the orthosteric binding site (OBS), hereafter termed as the extracellular offsite or EXO (Fig. 1d-e). This secondary region encompasses vestibular binding sites previously identified in A_2A_[34–36] as well as in other class A GPCRs,[37– 42], including the muscarinic receptor M2R resolved by X-ray in complex with a secondary ligand bound in this region.[43] Increasing evidences have shown that this region can act as a metastable stopover[39, 40] or a check point[37, 41] that can influence ligand selectivity[42, 44], cooperative/competitive effects[38], and even residence times.[45] Our results indicate that when NECA occupied the EXO, ZMA exhibited an increased transition frequency into the OBS, enhancing its binding likelihood. In addition, the presence of either ligand in this region was also associated with a marked slowdown in ZMA’s unbinding kinetics, effectively increasing its residence time. Interestingly, this effect appeared to be unidirectional — ZMA did not similarly influence NECA’s binding or unbinding. This suggests a hybrid competition-cooperation mechanism wherein both ligands compete for the OBS while NECA simultaneously facilitates ZMA’s binding — a phenomenon that, to our knowledge, has not been reported before. Finally, our protocol allows computing the ligands’ binding free energy and residence time, successfully reproducing the experimentally observed relative binding affinities of NECA and ZMA.

In the context of the growing focus on allosteric modulation and exosite targeting,[46–48] our approach establishes a foundation for elucidating the interplay between ligands at both shared and peripheral binding sites. By revealing how ligands influence one another’s binding pathway and residence time, it offers a new conceptual framework for drug design—one that seeks efficacy and selectivity by harnessing resilience to competition from endogenous ligands. In this light, multiligand simulations represent a significant technical advance, enabling a more mechanistically informed and precise approach to pharmacological innovation.

## 2 Results and Discussion

### 2.1 Ligand Competitive Binding Assays in A_2A_

For our in silico CBA, we constructed five simulation systems of the A_2A_ receptor embedded in a lipid bilayer, each containing 14 ligand molecules in varying NECA:ZMA ratios (Table 1). In particular, we investigated CBA in two *Pure Ligand Systems* — 100% NECA or 100% ZMA — and three *Mixed Ligand Systems* with differing relative NECA:ZMA concentration (29%:71%, 50%:50%, 71%:29%, respectively). All ligands were initially randomly placed on the extracellular side of the membrane and placed sufficiently far from the receptor to avoid introducing bias in diffusion or binding. A key bottleneck in unbiased multiligand simulations — and the primary reason an *in silico* CBA has remained elusive — is that unbound ligands tend to spend extended periods diffusing freely in the aqueous phase, with no guarantee of reaching the orthosteric binding site (OBS) within feasible simulation timescales. To overcome this, each system was simulated in 36 independent replicates of 30 *µ*s each, resulting in over 1 millisecond of aggregate CG simulation time per system (Table 1). Considering an acceleration factor of ~3 inherent to the Martini CGMD force field,[49] our CBA simulations effectively span seconds to minutes of real-world timescales— among the longest computationally accessible ligand–receptor binding windows reported to date— thereby capturing the most physiologically relevant binding dynamics.

**Table 1.**
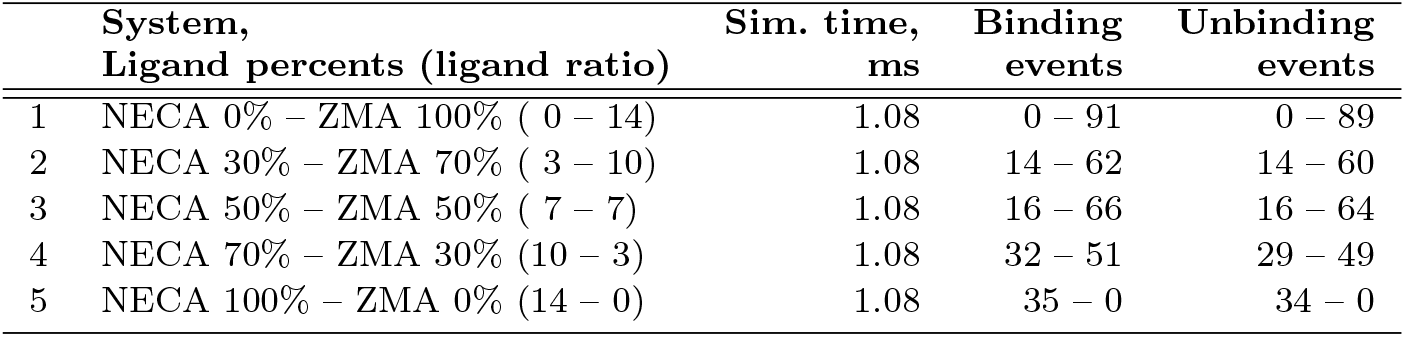
Systems investigated in the in silico CBA.

|  | System,<br>Ligand percents (ligand ratio) | Sim. time,<br>ms | Binding<br>events | Unbinding<br>events |
| --- | --- | --- | --- | --- |
| 1 | NECA 0% – ZMA 100% ( 0 – 14) | 1.08 | 0 – 91 | 0 – 89 |
| 2 | NECA 30% – ZMA 70% ( 3 – 10) | 1.08 | 14 – 62 | 14 – 60 |
| 3 | NECA 50% – ZMA 50% ( 7 – 7) | 1.08 | 16 – 66 | 16 – 64 |
| 4 | NECA 70% – ZMA 30% (10 – 3) | 1.08 | 32 – 51 | 29 – 49 |
| 5 | NECA 100% – ZMA 0% (14 – 0) | 1.08 | 35 – 0 | 34 – 0 |

Since the free ligands had no prior “knowledge” of the binding site, they needed to “discover” the orthosteric binding conformation themselves. Remarkably, our CGMD protocol successfully reproduced poses matching the experimentally determined binding modes of both NECA and ZMA. We observed lowest RMSD values of *~*0.2 nm and *~*0.3 nm, respectively, relative to their crystallographic poses in A_2A_ (PDB IDs 2YDV[50] and 3EML[51]) (Fig. 2). The observed broad RMSD distributions is consistent with the expected variability of ligand conformations at the CG resolution,[49] including ligand longitudinal inversion, angular deviations, and lateral shifts within the binding pocket.

**Fig. 2.**
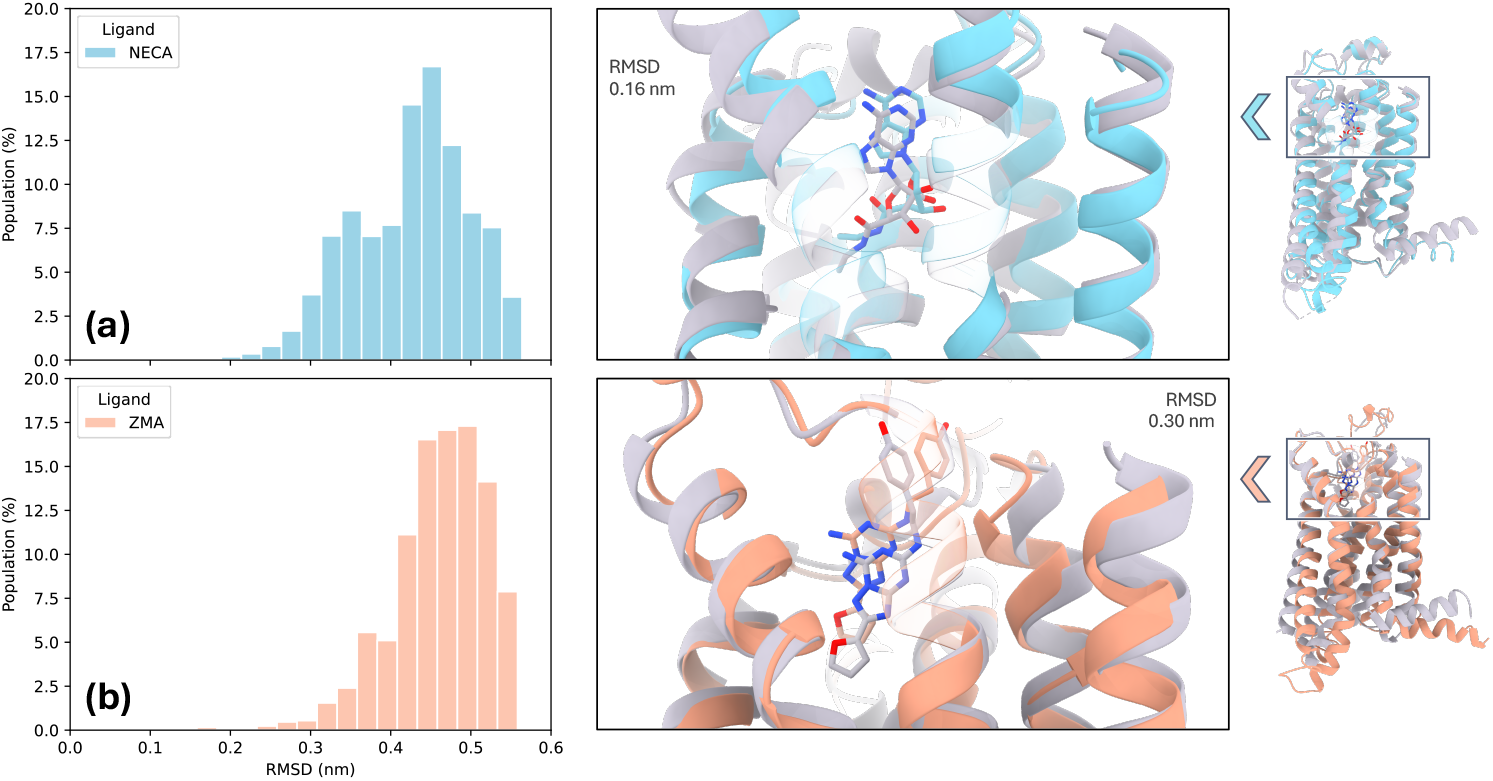
Orthosteric binding poses of NECA and ZMA from CG simulations. **(a-b)** RMSD distributions of NECA and ZMA binding poses from CG simulations, computed relative to the crystallographic references 2YDV and 3EML, respectively. To the right of each, a representative low RMSD pose is shown, backmapped to atomistic resolution. For clarity, RMSD distributions of OBS poses within 0.55 nm of the experimental structures are shown, and portions of TM6 and TM7 are rendered with increased opacity.

While the OBS is central to the pharmacology of ligands like NECA and ZMA, other sites along the ligand entry path can profoundly influence binding dynamics. In our simulations, ligands were free to diffuse across the solvent, the lipid bilayer, and the protein surface. By monitoring positional stability and motility, we identified a distinct region preceding the OBS where ligands frequently paused or interacted. This secondary region, confined primarily by the second extracellular loop (ECL2), is hereafter referred to as the *EXO* (Fig. 1d-e). The EXO includes intermediate ligand binding sites and vestibular pockets identified in previous GPCR studies, such as regions directly above the OBS, enclosed by ECL2 and ECL3, and adjacent to transmembrane helices TM6 and TM7 (Fig. S1). [34–43]. Its spatial location and interaction patterns suggest that it functions as an antechamber for ligand entry, where competition or cooperation among ligands can emerge before reaching the OBS. To more accurately capture ligand behavior as it transitions from free diffusion to protein engagement, we also defined an extended *Pre-Protein Environment* (PPE), encompassing the solvent-accessible region immediately adjacent to the receptor’s extracellular surface. (Fig. 1d-e). This framework allowed us to analyze ligand-ligand and ligand-protein interactions as a continuum from fully solvated states to productive binding, revealing how early-stage interactions can influence downstream occupancy and dynamics at the OBS.

### 2.2 Ligand binding mechanism

In our CBA, we define a successful binding event as a complete cycle comprising two distinct phases: the *binding phase* and the *unbinding phase*. The binding phase involves the ligand’s diffusion from the bulk solvent to the OBS, while the unbinding phase constitutes the ligand’s dissociation from the OBS and its return to the solvent.

Before reaching the OBS, ligands frequently interact with the intermediate regions we have designated as the EXO and the PPE. These peripheral zones exhibit transient ligand interactions compared to the OBS, as indicated by shorter binding and dissociation times (Fig. S2; see also next section for detailed kinetics). Notably, only a subset of ligands that bind to the EXO eventually reach the OBS, as shown by the computed number of binding events to the OBS for the ligands staying at the EXO (Table S1–S2). This observation supports the role of the EXO as a dynamic intermediate zone — a transient staging ground where ligands may bind temporarily, encounter other ligands, and engage in competitive or cooperative interactions, before eventually moving to the OBS.

Initially, the receptor is unoccupied, with multiple ligands simultaneously competing for access to the same binding site. Within the defined spatial hierarchy of PPE, EXO, and OBS, this gives rise to various co-occupancy scenarios in which two or more ligands simultaneously engage different regions of the receptor (Fig. 3a). Of particular interest are the decisive cases where a ligand in the EXO transitions to the OBS while another ligand simultaneously occupies the EXO or nearby, highlighting direct competitive dynamics and revealing potential facilitative mechanisms. As shown in Fig. 3b-c, ZMA transitions to the OBS occur more frequently in the presence of NECA than vice versa, indicating that *when both ligands are present, ZMA is the typical winner of competitive binding*. Notably, these decisive events predominantly occur in the EXO, reinforcing the region’s role as a critical mediator in ligand selection and entry. Intriguingly, although NECA is less dominant in the direct competition for the OBS, it exerts a clear facilitative effect on ZMA’s binding. Specifically, increasing NECA concentration correlates to a marked increase in ZMA transitions to the OBS (Fig. 3b). Conversely, ZMA does not play any effect on the binding of NECA (Fig. 3c). To ensure fair comparisons across systems with varying ligand abundance, all transition counts were weighted to account for differences in ligand concentration (see Methods, Section 4.6). These findings uncover a previously unreported cooperative mechanism in which NECA enhances ZMA binding, revealing a cooperative dimension within a system traditionally viewed as a purely competitive process.

**Fig. 3.**
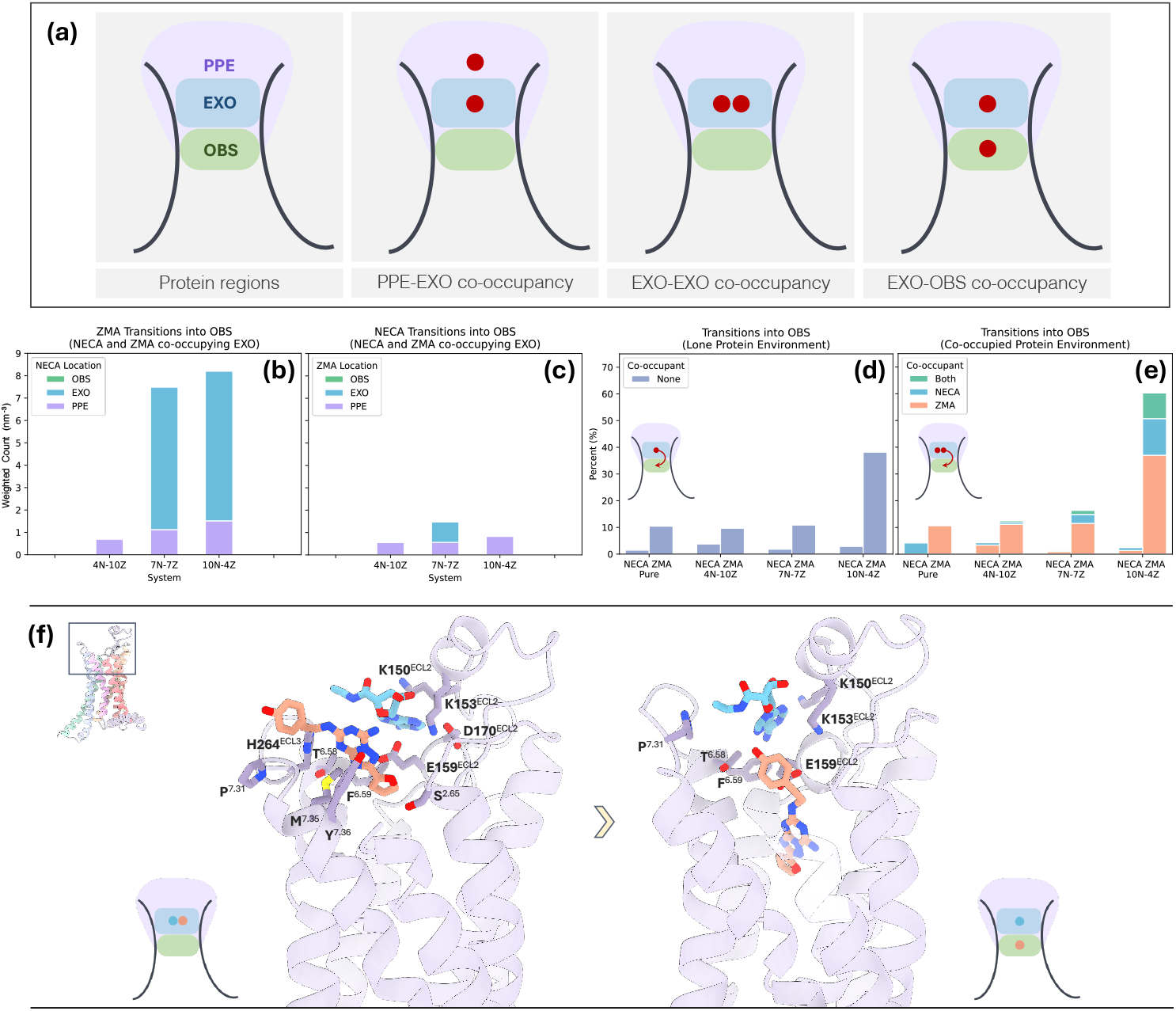
Protein co-occupancy reveals signatures of competition and cooperation. **(a)** Schematic of the receptor regions, illustrating three representative cases of co-occupancy by two ligands. **(b-c)** Frequency of successful EXO-to-OBS transitions in the presence of a second ligand, colored according to the region occupied by the co-occupant. **(d)** Fraction of EXO-to-OBS transitions in the absence of a co-occupying ligand ligand. **(e)** Fraction of EXO-to-OBS transitions in the presence of a co-occupying ligand at the EXO, colored according to the identity of the co-occupant. **(f)** Atomistic backmapped structures illustrating competitive and cooperative binding mechanisms of NECA and ZMA at the OBS. *Left)* NECA and ZMA co-occupying the EXO site formed by ECL2, ECL3, TM6, and TM7, prior to ZMA transitioning into the OBS. *Right)* Final binding mode of ZMA at the OBS in the presence of NECA at the EXO. CG structures corresponding to ZMA binding at the OBS with the lowest RMSD relative to the experimental binding pose were selected for atomistic backmapping.

Intrigued by these observations, we performed a more detailed analysis of ligand transitions. Specifically, we examined all transitions originating from the EXO region, quantifying the fraction that successfully reached the OBS, and distinguishing between events occurring in a ligand-free protein and those involving protein co-occupancy (Fig. 3d-e). This comparison revealed clear differences in transition behavior between these two environments. Particularly, when NECA was present in the EXO, the rate of ZMA transitions to the OBS increased markedly (blue and green histograms, Fig. 3e). This effect scaled with NECA concentration, underscoring the specificity and dose-dependence of the cooperative interaction. In contrast, ZMA transitions were virtually identical in ligand-free settings and when co-occupied by another ZMA molecule (Fig. 3d and pink histograms in Fig. 3e), suggesting that cooccupancy by a homologous ligand exerts no notable mechanistic influence.

Together, these results reveal a nuanced interplay between competitive and cooperative binding dynamics. Traditional models of ligand competition focus on direct displacement at the OBS. Our results indicate that such competition for the OBS may be preconditioned by earlier events at peripheral binding regions. In the A_2A_ case, the EXO acts as a mechanistic gatekeeper: cooccupancy at this site biases the competitive outcome in favor of one ligand (ZMA) over the other (NECA), even before the OBS is engaged. At the EXO site, both NECA and ZMA engage residues on extracellular loops 2 and 3, including K150^ECL2^, K153^ECL2^, E169^ECL2^, and H264^ECL3^ (Fig. 3f). Notably, E169^ECL2^, and H264^ECL3^ were recently identified by NMR studies of wildtype and mutant A_2A_ as key determinants of ZMA’s residence time.[52] The A_2A_ EXO spatially overlaps with the intermediate or vestibular binding sites reported in other class-A GPCRs, where such regions have been implicated in modulating ligand binding. [34–36, 53, 54] This correspondence underscores the EXO as a potentially generalizable modulatory element of GPCRs and highlights the importance of further exploring its structural and functional roles across this receptor family.

### 2.3 Ligand residence times in A_2A_

In our CBA simulations, we observed a remarkable number of spontaneous ligand binding and unbinding events (Table 1). During the *Unbinding Phase*, approximately 50% of ligand dissociation events from the OBS occurred while the EXO region was simultaneously occupied (Fig. S3b). This is markedly higher than the ~20% co-occupancy observed during the Binding Phase (Fig. S3a). Given that the only viable exit pathway from the OBS passes through the EXO, these findings highlight the need to understand how EXO occupancy influences ligand unbinding from the OBS. Specifically, the *Unbinding Phase* is defined as the time interval during which a ligand leaves the OBS and reaches the fully solvated state - an event that determines the ligand’s residence time (RT) within the receptor. Ligand dissociation from the receptor typically follows a Poisson process (see Methods 4.7 and ref [55]), such that, with adequate sampling and a sufficient number of binding–unbinding events, the distribution of unbinding times can be modeled as an exponential function with a rate parameter *λ* = 1*/τ*, where *τ* represents the ligand’s RT. As a key kinetic descriptor of the unbinding process, the RT provides a direct and quantitative means to compare ligand behavior under conditions of single occupancy versus receptor co-occupancy.

However, quantifying the specific influence of EXO-region occupancy on ligand unbinding from the OBS presented several challenges. First, in multiligand simulations involving several copies of each ligand type, interactions between EXO–OBS ligand pairs and other free ligands may occur, introducing potential confounding factors that are difficult to disentangle. Second, EXO binding is more transient than OBS occupancy (Fig. S2), often allowing multiple, sequential EXO binding events to occur during a single OBS residence interval. This dynamic behavior complicates the assignment of causality, especially in mixed-ligand environments where the identity and timing of EXO occupants vary over the course of a single OBS-binding episode.

To enable a focused investigation of the influence of an EXO-bound ligand on the unbinding of an OBS-bound ligand, we isolated OBS-EXO ligand pairs into environments containing only the two. Specifically, we extracted frames of stable EXO–OBS co-occupancy from the original CBA simulations, covering all possible OBS-EXO ligand pairings (Fig. 4 central graphics). From each selected frame, we set up new simulation systems with only the OBS and EXO ligands, preserved their poses during equilibration, and then ran unbiased production MD calculations. As a *control*, we generated analogous systems in which the EXO ligand was removed from the same starting frame, isolating the OBS-bound ligand under otherwise identical conditions. Each system was subjected to at least 100 independent simulation replicas to ensure robust statistical sampling. Each simulation was terminated on the dissociation of the OBS ligand from the receptor, with the elapsed simulation time recorded as the unbinding time for that replica. For additional comparison, we also computed RTs for NECA and ZMA directly from the original CBA trajectories in the pure systems. The results are presented in the Supplementary Information (Supplementary Discussion S3.1, Fig. S4-S5 and Table S3).

**Fig. 4.**
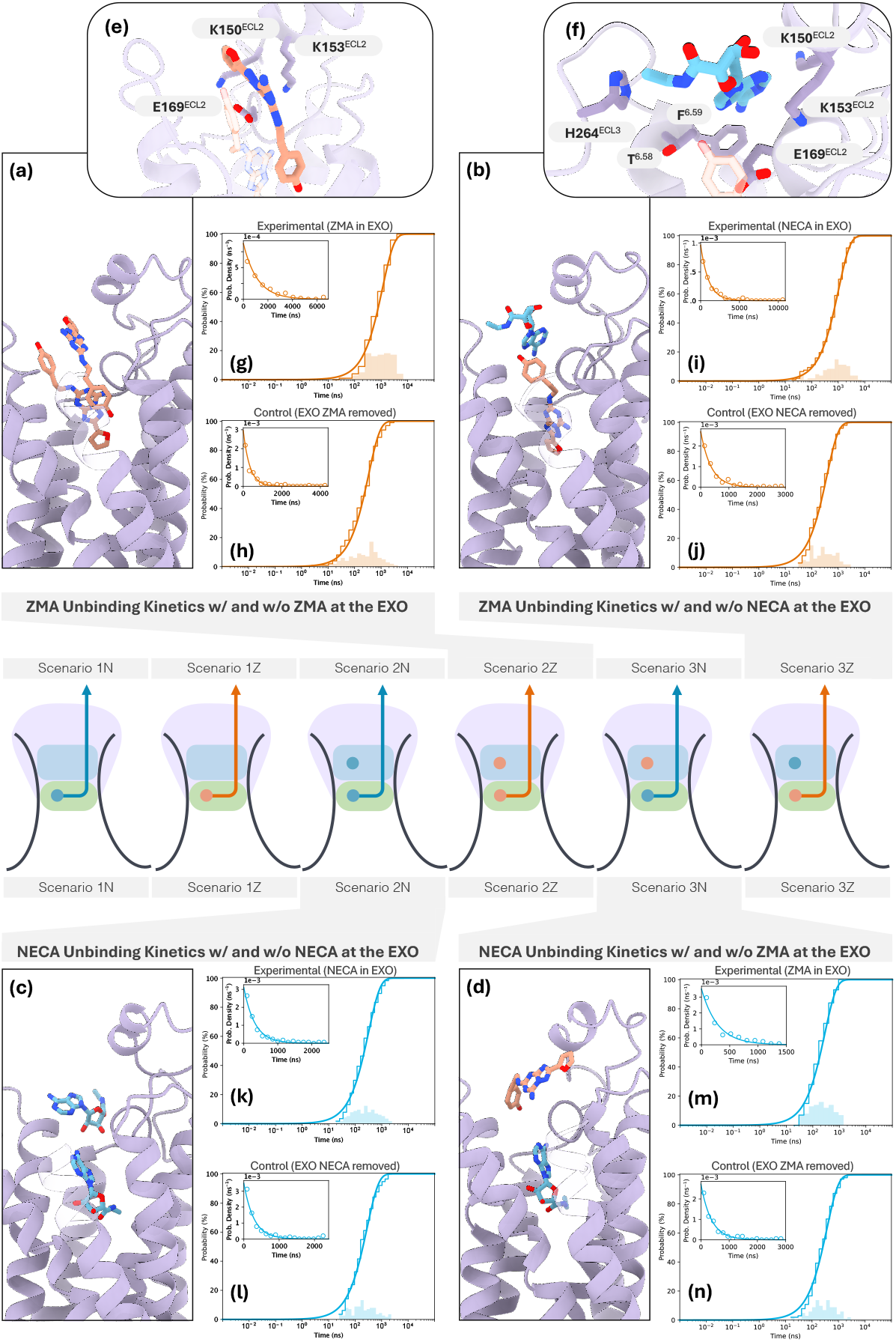
Protein co-occupancy and its influence on unbinding kinetics. The central schematic illustrates different unbinding scenarios for NECA (blue) and ZMA (orange). **(a-d)** Representative atomistic structures (backmapped) from unbinding simulations, showing EXO–OBS co-occupancy. In these trajectories, the EXO ligand typically dissociates first, while the OBS ligand remains bound for a longer duration. **(e-f)** Zoomed views of the receptor cooccupied by EXO ligands and ZMA in the OBS, highlighting receptor residues available for interaction. Further details and cluster analysis of EXO poses are provided in Supplementary Information (Section S3.2). **(g-n)** Poisson process modeling of OBS ligand unbinding. Each panel pair compares unbinding simulations with an initial EXO ligand (top) to control systems without the EXO ligand (bottom). Foreground curves show cumulative histograms of unbinding times (ragged line) fitted with the theoretical cumulative distribution function (CDF) (solid line), where the *y*-axis represents the probability *P*_*n*=1_(*t*) that the ligand has dissociated by time *t*. Background histograms display the same data non-cumulatively on a logarithmic time scale, with the *y*-axis representing the probability *P* (*a* ≤ *t* ≤ *b*) of a dissociation event occurring within the time interval [a,b]. Insets show empirical (o) and theoretical (solid line) exponential probability density functions of transition times.

In Fig. 4, we present the RT probability density functions for all the mixed-ligand systems, namely, NECA at the OBS with NECA or ZMA at the EXO, and ZMA at the OBS with ZMA or NECA at the EXO. For comparison, we also report the RT probability density functions for the single-ligand occupancy (*Control* in Fig. 4). The main plots display the empirical cumulative distribution function (eCDF) of unbinding times, representing the observed probability *P* (*t*) that the ligand dissociates by time *t*. These curves were fitted using Eq. 6, the theoretical cumulative distribution function (tCDF) of a Poisson process, from which the RT *τ* was extracted (see Methods 4.7 for details).

The RT values for all unbinding scenarios are reported in Table 2, together with three statistical indicators used to assess data reliability: (1) the *p*-value from a two-sample Kolmogorov-Smirnov (KS) test;[56] (2) the ratio *m*_*t*_*/m* between the theoretical median *m*_*t*_ and the empirical median *m* calculated from the fit of tCDF and eCDF; and (3) the ratio *µ/σ* of the empirical mean to its standard deviation. All systems passed the KS test with *p*-values above the 0.05 threshold, indicating a statistically significant agreement between the tCDF and the eCDF. The *m*_*t*_*/m* ratio provides an additional parametric validation: since the median is less sensitive to long-tailed distributions and outliers, a ratio close to unity supports the model’s accuracy in reproducing the theoretical kinetic data. Our results exhibit excellent consistency, with most *m*_*t*_*/m* value near 1. Similarly, the observed *µ/σ* ratios are reasonably close to unity, as expected for exponentially distributed data. Together, these indicators confirm that the unbinding times follow an exponential distribution — consistent with a Poisson process— thereby validating the statistical robustness of our RT estimates. It should be noted, however, that RTs obtained from CG simulations are systematically shorter than those measured experimentally or from atomistic simulations, owing to the intrinsic acceleration of molecular dynamics at the coarse-grained resolution.[29] This acceleration factor is inherently system-dependent, reflecting the degree of free-energy surface smoothing introduced by coarse-graining. Nevertheless, given the high structural and compositional similarity among our simulated systems, we can reasonably assume comparable acceleration factors across all cases, enabling meaningful relative comparisons of RTs between different ligand pairings.

**Table 2.**
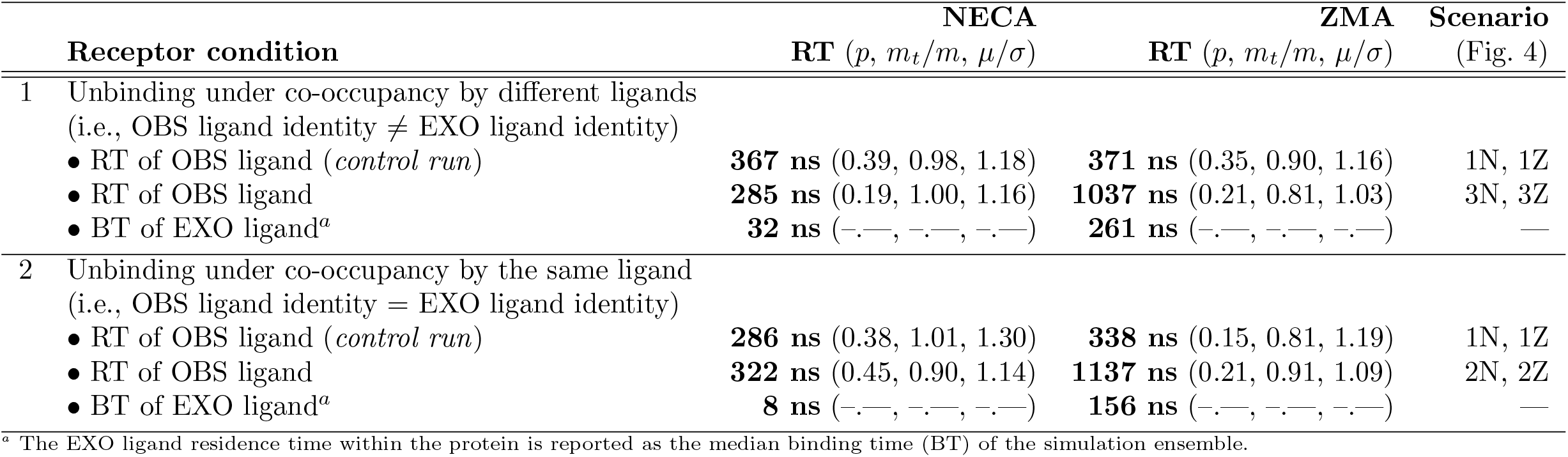
Ligand Residence Times (RT) of NECA and ZMA under single- and co-occupancy conditions.

As notably shown in Table 2, when the EXO is occupied, ZMA dissociation from the OBS is markedly slowed, showing an approximately threefold increase in RT. Unlike ZMA association — which displayed ligand-dependent sensitivity — the unbinding slowdown appears ligand-insensitive, occurring irrespective of whether NECA or ZMA occupies the EXO site. In contrast, NECA unbinding from the OBS remains unaffected by EXO occupancy, yielding comparable RTs whether the EXO is vacant or co-occupied, and regardless of the co-occupant’s identity. Furthermore, the RT of ligands bound at the EXO is significantly prolonged when ZMA occupies the OBS — regardless of the EXO ligand’s identity — with RTs increasing by a factor of 8 to 20 relative to systems in which NECA occupies the OBS (see BT of EXO ligand in Table 2). In other words, the ligands exert a reciprocal temporal effect, indicative of a cooperative interaction that is asymmetric with respect to ligand identity. Specifically, OBS-bound ZMA and any EXO ligand mutually prolong each other’s binding times, whereas OBS-bound NECA exerts no measurable influence on the RT of co-occupying EXO ligands. Our results point to a central role of EXO-site residues such as K150^ECL2^, K153^ECL2^, E169^ECL2^, and H264^ECL3^, in shaping ligand residence times. These residues, which mediate ligand–EXO interactions (Fig. 4e–f and Supplementary Discussion S3.2 for details), are poorly conserved across adenosine receptor subtypes (A_1_, A_2A_, A_2B_, A_3_), potentially contributing to subtype binding selectivity. Together, these findings highlight the pivotal role of the EXO region in modulating ligand binding kinetics and reveal a cooperative mechanism between ligands that governs the duration of OBS occupancy, consequently influencing downstream receptor signaling dynamics.

### 2.4 Ligand/A_2A_ binding free energy

As the final component of our CBA protocol, we evaluated ligand binding affinity. While direct binding assays typically express affinity as a dissociation constant *K*_*d*_, computational studies more commonly report it as the standard binding free energy 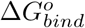, which is mathematically related to the association constant *K*_*a*_ or binding constant *K*_*bind*_, where *K*_*a*_ = 1/*K*_*d*_) (see Methods 3.8 and ref [18, 19] for details). Accurate and quantitatively converged estimates of 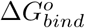 demand exhaustive sampling of both bound and unbound conformational ensembles.[57] Yet, conventional AA simulations fall short in this regard due to the rugged free-energy landscapes and high barriers typical of biomolecular systems, thereby requiring the use of advanced methodologies such as alchemical free-energy calculations or enhanced-sampling techniques.[19, 58–60] This limitation can be overcome through CG representations, which reduce the number of degrees of freedom and thereby smooth the free-energy landscape. As a result, the system’s dynamics are accelerated, enabling statistically robust sampling of long-timescale processes such as ligand–protein association and dissociation.[29, 49]

In the case of A_2A_, which resides within the membrane plane, the ligand binding process can be described by a three-state equilibrium model involving the ligand (L), the protein (P), and the membrane (M):

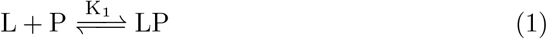

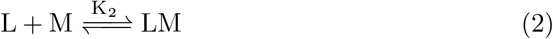

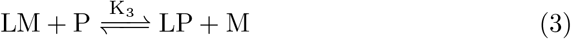

Fig. 5a provides a schematic of this three-state model. Inspired by protocols originally developed for estimating ligand binding affinities in unbiased atomistic simulations,[61–67] and recently adapted by our group for CG protein–ligand calculations,[49] we computed the absolute ligand/A_2A_ binding free energy 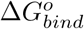 from the one-dimensional potential of mean force *PMF* (*r*).Because A_2A_ is membrane-embedded and not fully solvated, we restricted the integration volume to a spherical sector extending from the OBS toward the extracellular space, rather than a full spherical volume as in conventional protocols for soluble proteins (see Supplementary Discussion S3.2 for details). This ensures that the computed ligand concentration in bulk solution *L* accurately reflects equilibrium with both receptor-bound *LP* and membrane-associated *LM* states. Notably, the latter is strongly influenced by the ligand’s affinity for the phospholipid bilayer, a factor explicitly accounted for in our calculations (see a ligand parameterization feature detailed in Table S9).

**Fig. 5.**
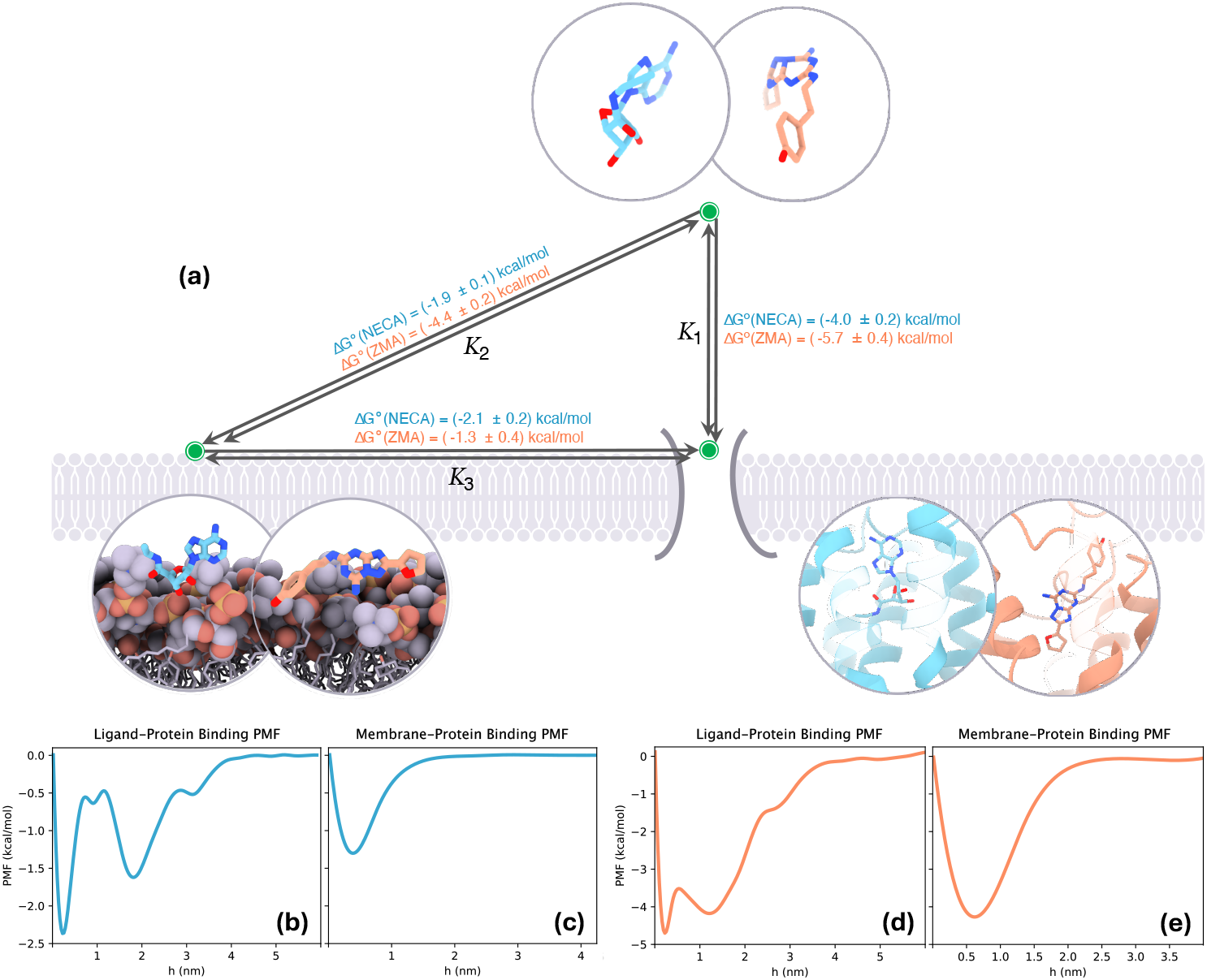
Thermodynamics of binding and competing equilibria. **(a)** Thermodynamic scheme illustrating the three co-existing equilibria (bidirectional arrows) at the membrane–receptor _2A_ interface together with the corresponding standard Gibbs free energies calculated from CG simulations. **(b-e)** Potentials of mean force (PMFs) describing NECA and ZMA binding to A_2A_ and their partitioning into the membrane.

Specifically, the ligand/A_2A_ binding PMFs for NECA and ZMA were calculated as the average free-energy profiles along the dissociation coordinate, spanning from the fully bound state *LP* at *r* = 0 to the unbound state *L* at *r* ≥ *R*_*cutoff*_ (Fig. 5b and Fig. 5d). Analogously, ligand–membrane PMFs were obtained by tracing the free-energy landscape from the membrane-bound *LM* state at *r* = 0 to the unbound state *L* at *r ≥ R*_*cutoff*_ (Fig. 5c and Fig. 5e). The resulting 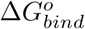 values, reported in Fig. 5 and Supplementary Table S4, correspond to the free-energy differences between the bound and unbound state, computed using data from the *Pure Systems* to avoid cross-ligand interference (see Supplementary Table S5 for 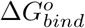 values in the *Mixed Systems*). Our results indicate that ZMA binds more strongly to A_2A_ than NECA, with a 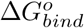 1.7 kcal*·*mol^*−*1^ lower than NECA (−5.7 *±* 0.4 kcal*·*mol^*−*1^ and −4.0 *±* 0.2 kcal·mol^*−*1^, respectively), in line with the ~1.8 kcal·mol^*−*1^ difference observed experimentally ([−12.5, −12.1] kcal·mol^*−*1^ for ZMA vs. [−11.9, −9.5] kcal·mol^*−*1^ for NECA). The absolute values are underestimated, likely due to the Martini 3 CG force field overestimating hydrophilic interactions and thus favoring the solvent-exposed unbound state relative to the more hydrophobic protein-bound state.[68] Nevertheless, given the structural and compositional similarity across systems, relative comparisons remain meaningful. ZMA also exhibits higher membrane affinity than NECA, with computed 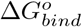 values of −4.4 *±* 0.2 kcal·mol^*−*1^ and −1.9 ±0.1 kcal·mol^*−*1^, respectively. Notably, for ZMA, membrane partitioning is of comparable magnitude to its receptor binding free energy, indicating that membrane association constitutes an energetically significant competitive process for receptor binding, as described by Eq. (1) and Eq. (2). This highlights a documented but often overlooked phenomenon: aqueous ligands partitioning into the lipid bilayer can modulate apparent receptor affinity, with important implications for drug discovery and design. [69]. Finally, the derived binding free energies 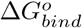 for *LP* association via lateral diffusion from the membrane to the protein (Eq. (3)), indicate an energetically disfavored pathway. Nonetheless, such membrane-mediated binding routes have been previously observed and remain of mechanistic interest, particularly for understanding alternative ligand entry pathways and transient binding intermediates.[70]

## 3 Conclusion

In this work, we introduce an in silico Competitive Binding Assay (CBA) that captures the full complexity of multiligand receptor engagement under near-physiological conditions. Using the A_2A_ adenosine receptor with its agonist NECA and inverse agonist ZMA as a model system, our approach simulates spontaneous diffusion of both ligands between the receptor, surrounding solvent, and membrane interface, reconstructing association and dissociation events in the presence of competing ligands. This yields direct structural, kinetic, and thermodynamic insight into cooperative and competitive binding mechanisms. Without prior structural information, both ligands reproduced their experimental orthosteric poses and revealed an intermediate binding region (EXO) preceding the OBS, delineated by ECL2, ECL3, and adiacent TM helices. This vestibular domain - corresponding to previously reported metastable binding sites across class A GPCRs - can accommodate multiple ligands simultaneously, acting as a dynamic arena for ligand–ligand interactions and kinetic modulation. Within this region, NECA and ZMA directly compete for access to the OBS, with ZMA consistently outperforming NECA. Strikingly, this advantage was amplified by the presence of NECA, unveiling a previously unrecognized cooperative effect. The facilitation of ZMA’s orthosteric transitions by NECA was concentration-dependent (i.e., it is enhanced by increasing the NECA concentration), highlighting a cooperative–competitive interplay within a system traditionally regarded as purely competitive. Notably, this effect was asymmetric: NECA’s binding success remained unchanged by ZMA’s presence, revealing a directional specificity in the underlying mechanism.

In our multiligand simulations, which closely mimic the crowded and dynamic physiological milieu, ligand dissociation acquires an additional layer of complexity: the influence of a second ligand transiently occupying the receptor’s vestibular region. We observed a striking correlation between the residence time (RT) of a primary orthosteric binder and the presence of a secondary ligand at the EXO. This relationship is not merely binary but directionally specific: ZMA’s RT at the orthosteric site is markedly prolonged when either NECA or ZMA occupies the EXO, whereas NECA’s RT remains unaffected by any EXO-bound ligand. This asymmetry reveals a form of ligand-specific “cognizance” at the EXO, which works as an active modulatory site capable of influencing ligand access, retention, and egress in a ligand-dependent manner. The concept resonates with emerging paradigms in GPCR pharmacology, where allosteric modulators such as PAMs, NAMs, and BAMs, fine-tune receptor behavior through transient or stable engagement at non-orthosteric sites.[16, 42**? ?**] The EXO thus emerges as a functional analog of such allosteric hotspots, modulating binding kinetics without directly overlapping with the primary binding pocket. Importantly, the EXO is delineated by residues K150_ECL2_, K153_ECL2_, E169_ECL2_, and H264_ECL3_, which differ markedly between A_2A_ and other adenosine receptors (e.g., A_2B_), offering a molecular basis for subtype selectivity. This insight introduces a new paradigm for tuning drug residence time through indirect modulation of orthosteric kinetics via EXO engagement. The pharmacological implications are significant such as designing bitopic ligands that simultaneously target the orthosteric and EXO regions with chemically distinct moieties, enabling fine control over residence time, receptor subtype selectivity, and functional outcome. Similar dual-site strategies have recently proven successful for the *µ*-opioid receptor, where bitopic fentanyl derivatives engaging both the OBS and the allosteric sodium-binding pocket, yielded compounds with improved analgesic and toxicity profiles.[71]

Our CBA also reproduced experimental trends in binding thermodynamics, yielding a 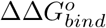 consistent with reported data. Notably, the computed membrane binding affinity of ZMA was comparable in magnitude to its receptor binding free energy, identifying the lipid bilayer as a major competitor to the receptor for ligand interaction. This finding emphasizes the necessity of explicitly accounting for ligand partitioning into the membrane, a highly nontrivial energetic process in which ligands, lipids, and proteins engage in simultaneous and interdependent binding equilibria. By disentangling these contributions, our approach provides a holistic picture of receptor–ligand binding, encompassing both direct protein engagement and membrane competition. Continued improvements in CG force-field accuracy are expected to further enhance the predictive power of in silico CBA, particularly in capturing the subtle energetic and solvation effects that govern binding thermodynamics.[68, 72, 73]

The present CBA framework transcends the classical 1:1 protein–ligand paradigm by introducing a computational titration strategy that systematically varies ligand molar ratios, bridging the gap between atomistic simulations and experimental competitive binding assays. This methodological advance provides a predictive means to assess how drug candidates compete with endogenous ligands, a key determinant of *in vivo* pharmacological efficacy and receptor bias. Moreover, in silico CBA enables quantitative exploration of reciprocal effects between chemically distinct ligands engaging different receptor regions, such as orthosteric and allosteric sites. This is particularly relevant for GPCRs, where allosteric modulators — including PAMs, NAMs, BAMs, peptides, cholesterol, and membrane phospholipids — have emerged as critical regulators of receptor conformation and function.[16] By capturing these multi-ligand interactions under near-physiological conditions that reflect realistic ligand molar ratios, our approach provides unprecedented mechanistic insight into cooperative and competitive binding processes, revealing how distinct ligands dynamically modulate one another’s binding pathways, residence times, and receptor accessibility.

Altogether, our findings expand the framework of dynamic structure–activity relationships (dySAR) in ligand–GPCR interaction[16] explicitly integrating ligand interdependencies and the kinetic, thermodynamic, and structural determinants that govern binding across multiple receptor regions. The in silico CBA represents a general and scalable strategy to probe competitive, cooperative, and allosteric interactions under physiologically relevant conditions. By unifying these dynamic aspects within a single computational framework, the CBA opens new avenues for the rational design of orthosteric, allosteric, and bitopic ligands, and for accelerating mechanism-driven, structure-based drug discovery in GPCRs and other membrane-bound targets in a way not accessible before.

## 4 Methods

### 4.1 CG modeling and AA backmapping

The coarse-grained (CG) model of the A_2A_ receptor used in this work was the same as in our previous study and was prepared according to the Martini 3 force field (see ref. [49] for details). Ligand atomistic-to-coarse-grained (AA-to-CG) mappings were generated using the *CG Builder* web tool (https://jbarnoud.github.io/cgbuilder), following the mapping scheme shown in Supplementary Fig. S6. In Martini 3, mappings are defined with respect to the center of mass (COM) of each bead, including hydrogen atoms. Atomistic backmapping was performed using HEroBM, an equivariant graph neural network (GNN) backmapping algorithm recently developed in our laboratory.[74]

### 4.2 Molecular dynamics simulation setup and details

All simulations were performed using the GROMACS 2020.6 molecular dynamics (MD) package,[75] in combination with the Martini 3 CG force field and the ELNEDYN elastic network to preserve the receptor’s secondary structure.[76] In all simulation boxes (approximately ~13 nm in each dimension), the protein was embedded in a POPC bilayer using the *insane*.*py* script.[77] Free ligands were then added on the extracellular side using the `gmx insert-molecules` routine. Each system was solvated with CG water and neutralized by adding NaCl to a final concentration of 0.15 *mol · L*^*−*1^ using `gmx solvate and gmx genion.` The assembled systems were subjected to energy minimization followed by an NPT equilibration prior to the production phase. Detailed parameters of the system setup and simulation protocols are provided in Tables S6–S7. All structural visualizations and renderings were prepared using VMD[78] and ChimeraX.[79]

### 4.3 Ligand parameterization

Ligand parameterization followed the standard Martini 3 procedure: (1) AA-to-CG mapping was designed according to Martini mapping principles; (2) bead types were assigned to reflect the chemistry of the atoms represented within each bead (Fig. S6); (3) bonded interactions and (4) non-bonded parameters were defined based on atomistic reference simulations and further refined using solvent-accessible surface area (SASA) and octanol–water partition coefficient (log *P*) calculations (Table S8 and S9).[49, 76]

### 4.4 Definition of receptor binding regions

The most relevant receptor binding sites were identified based on the persistent residence of ligands within specific spatial regions over time. To this end, the motion and position of each ligand molecule were tracked throughout the simulations via its center of geometry (COG). A sliding time window of 200 frames (corresponding to 2 ns) was applied to identify ligand residence events in which the standard deviation of the COG position did not exceed 0.03 nm. For each ligand, all COG coordinates from these residence events were then combined and clustered using the DBSCAN algorithm, as implemented in the *scikit-learn* package in Python. This procedure revealed a large, well-defined cluster corresponding to the orthosteric binding site (OBS; green cloud in Fig. 1d), hereafter denoted as the set of points *L*_*OBS*_. Additional clusters located above the OBS were grouped and designated as *L*_*EXO*_ (blue cloud in Fig. 1d). Finally, the Pre-Protein Environment (PPE) region was defined as the space located up to 0.9 nm from the backbone beads of the upper half of the receptor’s transmembrane helices.

### 4.5 RMSD calculations

Ligand RMSD values relative to the experimental binding pose were calculated after mapping the atomistic crystal structure of the A_2A_-ligand complex onto the CG representation. Al structures were subsequently aligned using the backbone beads of the upper halves of the receptor’s transmembrane helices TM1-TM7.

### 4.6 Statistical weighting of multi-ligand binding ensembles

To account for i) the differing probabilities of regions with different volumes to host a ligand at any given time, and ii) the differing probabilities of mixed NECA–ZMA systems with varying composition ratios to produce specific ligand pairings within the binding site, all statistics associated with both regional occupancy and pair identity were normalized by a weighting factor *w*_*i*_ defined as

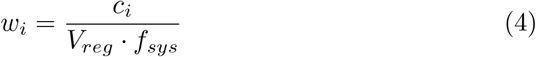

where *c*_*i*_ represents the raw statistic count, *V*_*reg*_ is the total volume of the regions occupied by a ligand pair *a* and *b*, and *f*_*sys*_ is a system-specific correction factor that reflects the relative probability of forming a specific ligand pair *a − b*. The value of *f*_*sys*_ was calculated as

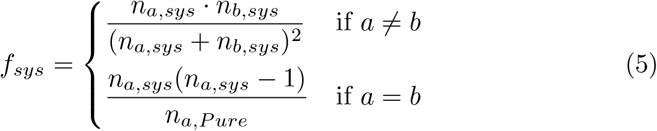

where *n*_*a,sys*_ and *n*_*b,sys*_ denote the numbers of ligand molecules of types *a* and *b* in any given system *sys*, and *n*_*a,P ure*_ is the number of ligand *a* molecules in the corresponding *Pure System* (reference system) containing only ligand *a*. This normalization ensures that occupancy and co-binding statistics account for both the accessible volume available for ligand binding and the expected combinatorial probabilities of ligand pair formation arising from the concentration effect in each mixed system.

### 4.7 Residence time calculations

If ligand dissociation events are independent, stochastic, and memoryless, tthey can be modeled as a Poisson process, where the time *t* to an unbinding event follows an exponential probability distribution *P* (*t*) = exp(−*t/τ*), with *τ* representing the mean residence time (RT). Accordingly, the probability *P*_*n*=1_(*t*) that a single unbinding event *n* = 1 has occurred by time *t* is given by the cumulative distribution function (CDF):

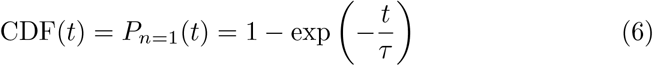

while the probability density of unbinding at time *t* is described by the probability density function (PDF):

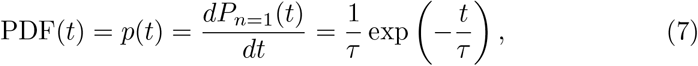

such that the integral of *p*(*t*) over any time interval [*a, b*] yields the probability *P* (*a* ≤ *t* ≤ *b*) that an unbinding event occurs between times *a* and *b*.

Residence times were obtained by fitting the observed unbinding times to Eq. 6 using the *curve fit* function from the *scipy* Python package. The resulting RT values are reported in Table 2. For all systems, the unbinding time was defined as the time interval between the ligand’s entry into the OBS and its subsequent crossing of the PPE boundary into the solvent, as illustrated by the top-left inset of Figs. S2c and S2f.

### 4.8 Free energy calculations

Let the point *C*_*LP*_ (*x*_*LP*_, *y*_*LP*_, *z*_*LP*_), corresponding to *r* = 0 of the ligand-protein (LP) interaction, denote the centroid of the ligand density within the orthosteric binding site (*L*_*OBS*_). *C*_*LP*_ is defined as the center of a sphere and the apex of a spherical sector of that sphere whose axis is aligned with the principal protein axis and has an aperture angle *α* (Fig. S7). An axial distribution function, ADF(*r*), was then obtained by binning the distances between *C*_*LP*_ and the subset of ligand positions located within the spherical sector, according to

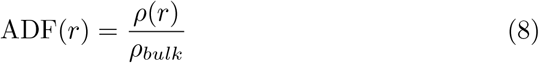

where *ρ*(*r*) = *n*(*r*)*/A*(*r*) is the local number density of ligand molecules, *ρ*_bulk_ = ∫_bulk_ *n*(*r*) *dr* ∫_bulk_ *A*(*r*) *dr* is the bulk number density, and ADF(*r*) therefore represents the ratio of the local to bulk ligand density, approaching unity at long distances. The surface area of the spherical cap corresponding to a given *r* is *A*(*r*) = 2*πr*^2^(1 *−* cos *α*). The bulk region was defined as extending from *R*_cutoff_ to *R*_max_, corresponding to the edge of the simulation box.

From ADF(*r*), the potential of mean force PMF(*r*) was computed as

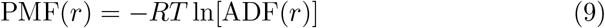

and, assuming that the LP interaction is approximately isotropic and radially symmetric, the binding constant was obtained by integrating

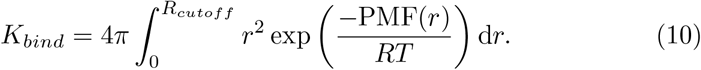

The corresponding standard binding free energy was then calculated as

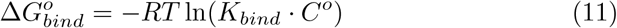

where *R* = 1.987 × 10^*−*3^ kcal·mol^*−*1^K^*−*1^, *T* = 310 K, and the standard concentration *C*^*o*^ = (1*/*1.66) nm^*−*3^. All parameters used are summarized in Supplementary Table S10. Relative to ref.[49], our method introduces the cone apex angle parameter *α*, whose value was determined based on the convergence behavior of the binding free energy calculations, as described in Fig. S8.

#### Calculation of ligand affinity to the membrane

Assuming the membrane lies parallel to the *xy*-plane, the *z*-axis aligned with the protein axis and orthogonal to the membrane (Fig. S9a). The center of the ligand-membrane (LM) interaction *C*_*LM*_ (*x*_*LM*_, *y*_*LM*_, *z*_*LM*_), corresponding to *r* = 0, was chosen such that *x*_*LM*_ and *y*_*LM*_ are positioned far from the protein as possible, in regions where the ligand distribution is not influenced by the protein and thus approximately uniform along the membrane plane. *z*_*LM*_ was defined as the centroid of the ligand’s spatial distribution along the *z*-axis. Using *C*_*LM*_ as the center of a sphere and as the apex of the spherical cone oriented along *z*, the radial distribution function (RDF), PMF, and 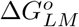 following the same procedure as for protein-ligand binding. The uncertainty in the calculation of 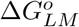 was estimated by evaluating the binding energy starting from various positions on the membrane plane, as described in Fig. S9b-c.

## Supporting Information

This paper comes with a *Supplementary Information* file in PDF format.

## Data Availability

Trajectories, structures, analyses, and all associated and auxiliary files are available upon request.

## Acknowledgments

This work has received funding from the European Research Council (ERC) under the European Union’s Horizon 2020 research and innovation programme (“CoMMBi” ERC grant agreement No.101001784), and it was supported by a grant from the Swiss National Supercomputing Centre (CSCS) under project ID s1293.

## Author contributions

V.L. designed and supervised the project. V.B.C performed the simulations and post-processing calculations. V.B.C., S.R., P.C., and V.L. analyzed the results and contributed to the writing of the manuscript.

