## Supplementary Information for "Beyond the 1:1 Ligand–Protein Paradigm: An In Silico Assay for Competitive Ligand Binding"

### Contents

**S1 Supplementary Figures** S2

**S2 Supplementary Tables** S8

**S3 Supplementary Discussion** S14

S3.1 NECA and ZMA residence time (RT) calculations from CBA simulations . . . S14

S3.2 EXO-OBS multiligand occupancy . . . S14

S3.3 Absolute ligand binding free energy calculation . . . S16

### S1 Supplementary Figures

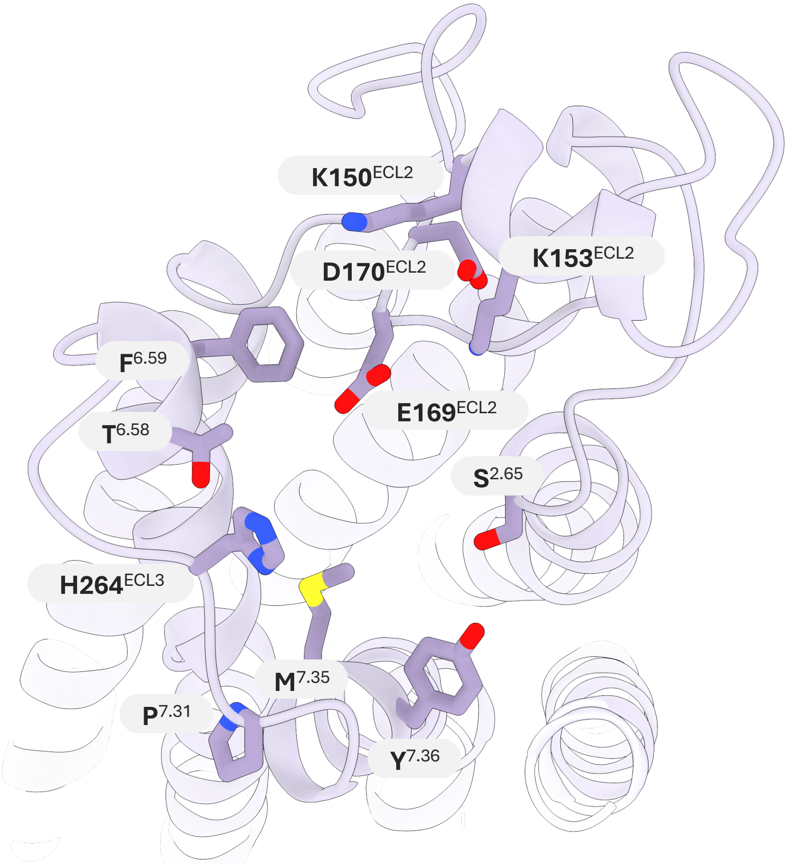


**Fig. S1 The EXO site of the $A_{2A}$ receptor** Residues defining the extracellular vestibular (EXO) site of the $A_{2A}$ receptor, labeled according to the Ballesteros–Weinstein numbering scheme. Located immediately above the orthosteric pocket, these residues from TM2, ECL2, TM6, ECL3, and TM7 form a transient interaction region where ligands can engage the receptor prior to orthosteric binding. This region serves as a dynamic interface supporting competitive and cooperative ligand interactions, as observed in our simulations.

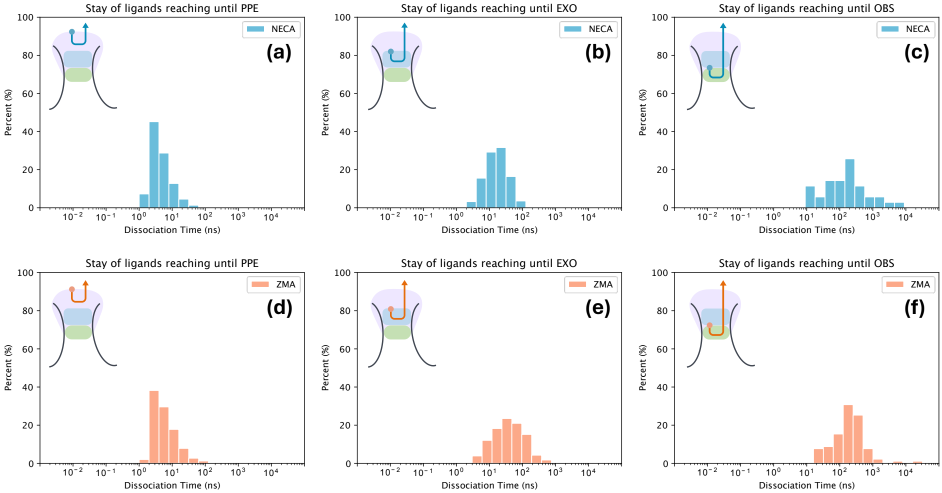


**Fig. S2** **Ligand dissociation time across protein regions** Distribution of ligand dissociation times from distinct receptor regions, computed from simulations of the *Pure Ligand Systems*. The $x$-axis represents the time required for a ligand to completely exit the protein after reaching a given region, as illustrated in the schematic at the top left of each panel. Panels **(c)** and **(f)** correspond to NECA and ZMA, respectively, and contain the same dissociation time data as in Fig. S5, but displayed here with a broader bin width for clarity.

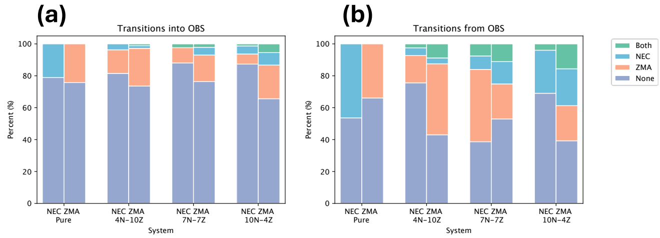


**Fig. S3** **EXO co-occupancy during ligand transitions at the orthosteric site** All observed transitions of ligands entering or leaving the OBS, color-coded according to the identity of the ligand simultaneously occupying the EXO at the moment of transition.

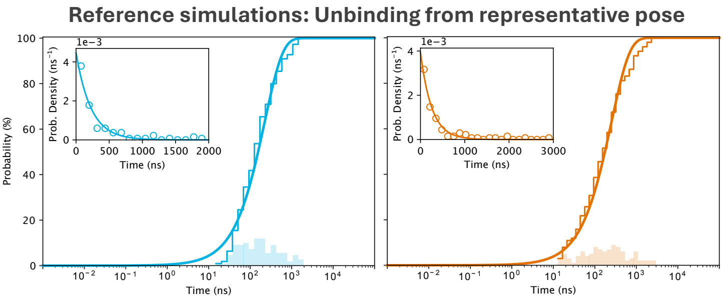


**Fig. S4 Poisson modeling of reference unbinding simulations** Unbinding time distributions for NECA (left) and ZMA (right) obtained from 100 unbinding simulations used to determine the residence time $\tau$ in a single OBS-ligand bound environment (R in Table S3). Each trajectory was initiated from a representative orthosteric pose corresponding to the centroid of the most populated cluster of OBS-bound conformations (RMSD $\leq$ 0.3 nm) identified in the original CBA simulations.

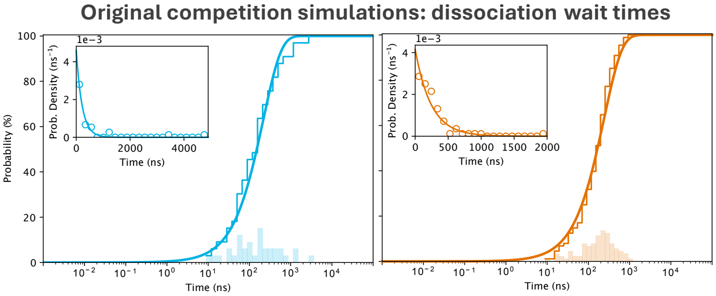


**Fig. S5 Poisson modeling of dissociation times from original CBA simulations** Distributions of orthosteric ligand dissociation unbinding times for NECA (left) and ZMA (right) obtained directly from the original CBA simulations of the *Pure Systems*.

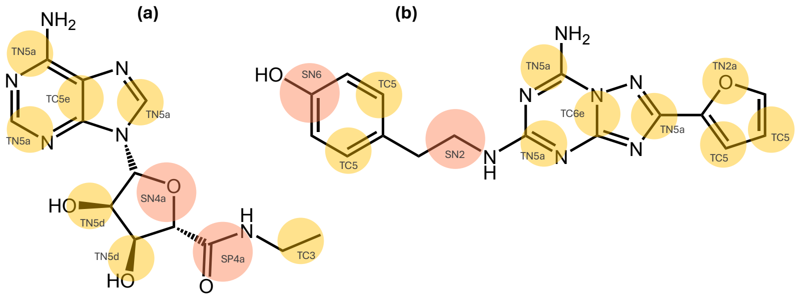


**Fig. S6 AA-to-CG ligand bead mapping for NECA and ZMA** CG representations of **a**, NECA and **b**, ZMA, constructed following standard Martini 3 protocols and best practices. The corresponding bead types are indicated, with beads color-coded by size: pink for small and yellow for tiny beads.

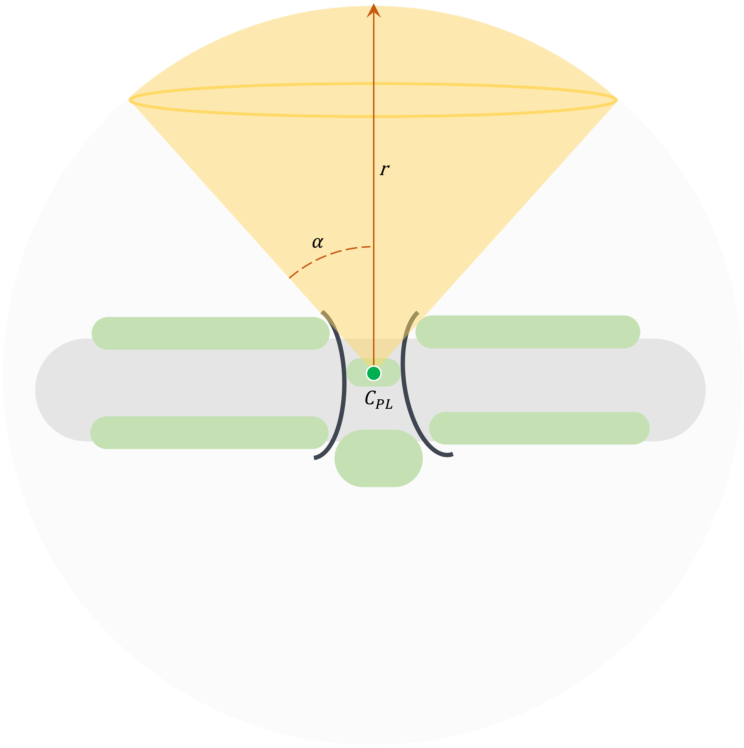


**Fig. S7 Spherical cone usage for the absolute ligand-receptor binding free energy** Schematic illustrating the selection of datapoints for computing the distribution function (DF), which is subsequently used to derive the PMF. The DF is calculated along the axis of a spherical cone extending from the observer (OBS, center, bound state) toward the solvent (edge, unbound state), where ligand-protein contacts are absent. The parameter $\alpha$ representing the cone's apex angle is indicated.

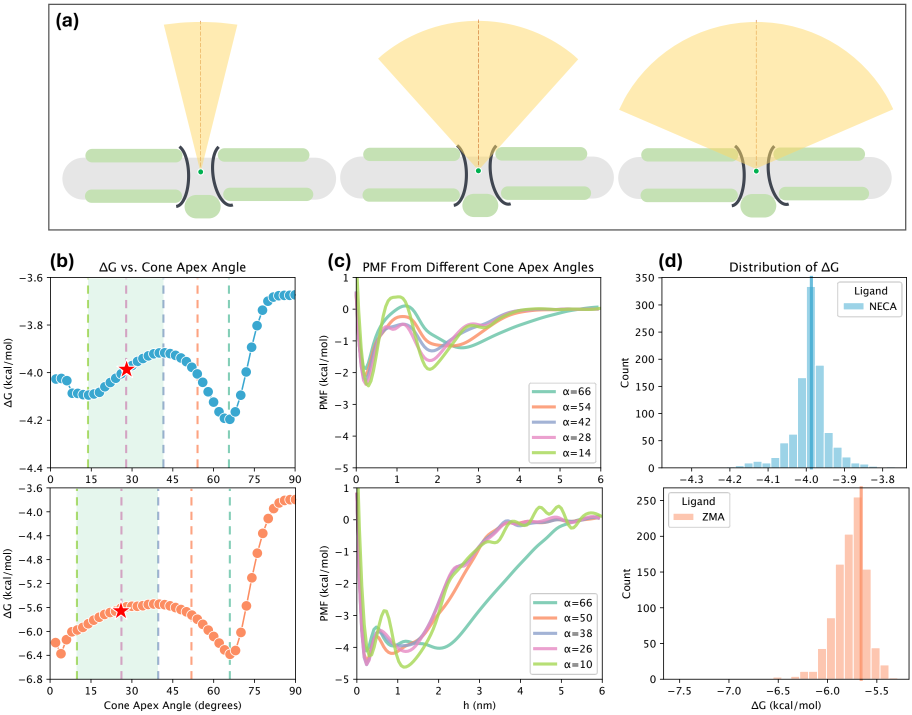


**Fig. S8 Selection of the apex angle $\alpha$ and associated uncertainty (a)** Illustration showing that overly small values of the conic aperture can exclude relevant protein volumes, whereas excessively large values can include ligand densities unrelated to protein-ligand interactions. **(b)** Evolution of the binding free energy as a function of the conic apex angle for NECA (top) and ZMA (bottom). As $\alpha$ increases from small values (approximately 20–40°), the computed free energy exhibits a slight increase due to the inclusion of ligand-inaccessible volumes occupied by protein side chains within the bound region. At larger $\alpha$ values, the free energy decreases as membrane-associated ligand densities begin to be included in the bound region. Beyond this minimum, the free energy rises sharply as membrane-ligand contributions are incorporated in both the bound and unbound regions. The most physically representative estimate of the ligand-receptor binding free energy lies in the range preceding the inclusion of inaccessible protein-occluded volumes. The reported value corresponds to the median across this interval (indicated by a star), and the span of this interval defines the associated uncertainty. **(c)** PMF profiles for NECA (top) and ZMA (bottom) at different apex angles. **(d)** Distribution of computed free energies obtained from random sampling of the unbound state using different values of the unbound density $\rho$ (Eq. 8 in the main text), serving as a sanity check since the ligand distribution function depends on $\rho$. Vertical lines (blue for NECA, orange for ZMA) indicate the binding free energy calculated using $\rho$ as defined by the spherical cone. The alignment of the vertical lines with the central region of the distributions, along with the overlap of the distributions with the reported uncertainty in *b)*, demonstrates consistency across calculations.

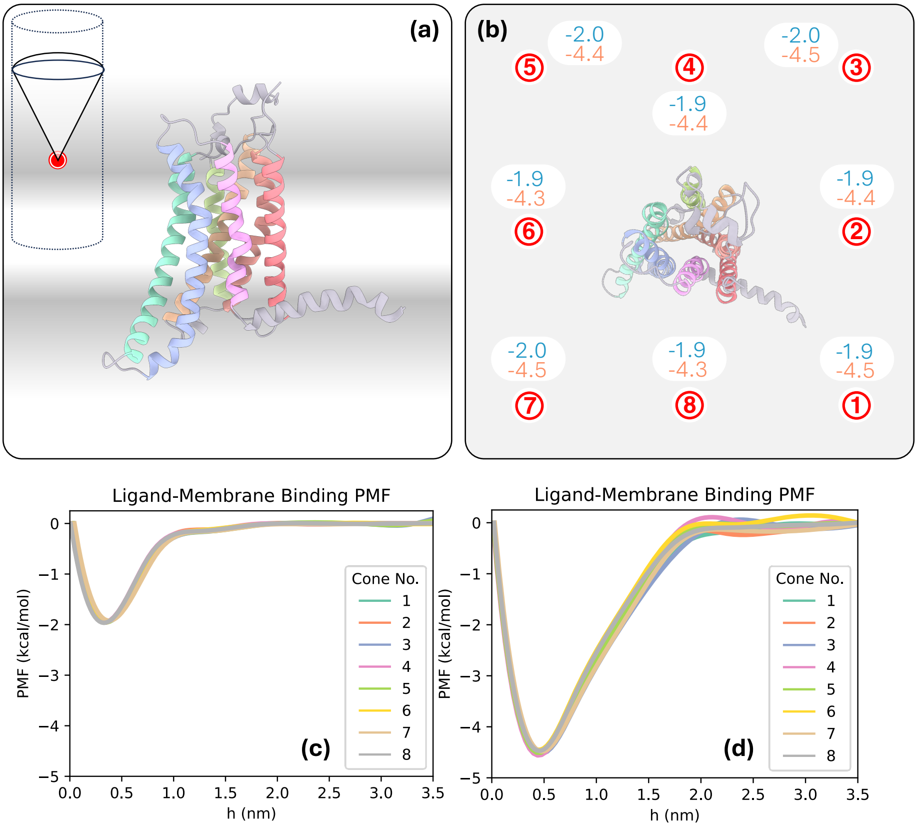


**Fig. S9 Calculation of ligand affinity to the membrane using the spherical cone approach (a)** The gray gradient schematically represents the ligand density observed in our simulations, highlighting the ligand–membrane interactions. To compute the $LM$ binding affinity, the apex of the spherical cone $C_{LM}(x_{LM}, y_{LM}, z_{LM})$ was positioned such that $z_{LM}$ corresponds to the $z$-coordinate of the center of mass of points within the smallest parallel cylinder that inscribes the cone. The $x_{LM}$ and $y_{LM}$ coordinates were chosen to maximize the distance from the protein, ensuring that the ligand positional distribution is minimally influenced by the protein and is as uniform as possible in the $xy$ plane. **(b)** (top view) Eight cones were placed in total (apices marked by red circles with "x") to verify the uniformity of ligand density far from the protein (cone parameters are detailed in Table S10). Free energy values are reported in $kcal \cdot mol^{-1}$, with NECA in blue and ZMA in orange. The mean values are reported in the main text, with uncertainties corresponding to the standard deviation. **(c-d)** PMFs generated from the eight samples for NECA and ZMA.

### S2 Supplementary Tables

**Table S1** Number of NECA binding events per region

| | **System** | *Events reaching* **PPE** | *Events reaching* **EXO**[a] | *Events reaching* **OBS**[b] |
|---|---|---|---|---|
| 1 | NECA 0% – ZMA 100% | - | - | - |
| 2 | NECA 30% – ZMA 70% | 9814 [100%] | 189 [ 2%] | 14 [ 7%] |
| 3 | NECA 50% – ZMA 50% | 18314 [100%] | 250 [ 1%] | 16 [ 6%] |
| 4 | NECA 70% – ZMA 30% | 14058 [100%] | 321 [ 1%] | 31 [10%] |
| 5 | NECA 100% – ZMA 0% | 32416 [100%] | 585 [ 2%] | 35 [ 6%] |

[a] Percentage of PPE events that ultimately reached the EXO region.
[b] Percentage of EXO events that ultimately reached the OBS region.

**Table S2** Number of ZMA binding events per region

| | **System** | *Events reaching* **PPE** | *Events reaching* **EXO**[a] | *Events reaching* **OBS**[b] |
|---|---|---|---|---|
| 1 | NECA 0% – ZMA 100% | 43202 [100%] | 606 [ 1%] | 91 [15%] |
| 2 | NECA 30% – ZMA 70% | 33449 [100%] | 1085 [ 3%] | 62 [ 6%] |
| 3 | NECA 50% – ZMA 50% | 26440 [100%] | 893 [ 3%] | 66 [ 7%] |
| 4 | NECA 70% – ZMA 30% | 12639 [100%] | 147 [ 1%] | 50 [34%] |
| 5 | NECA 100% – ZMA 0% | - | - | - |

[a] Percentage of PPE events that ultimately reached the EXO region.
[b] Percentage of EXO events that ultimately reached the OBS region.

**Table S3** Residence times (RT) of NECA and ZMA from original CBA simulations

| | **Simulation set** | **NECA** **RT** ($p$, $m_t/m$, $\mu/\sigma$) | **ZMA** **RT** ($p$, $m_t/m$, $\mu/\sigma$) | **Scenario** (Fig. 4) |
|---|---|---|---|---|
| R | Unbinding from a representative OBS pose in a singly occupied receptor (clustering of (OBS poses with RMSD $\leq$ 0.3 nm)[a] | **224 ns** (0.17, 1.02, 1.36) | **254 ns** (0.13, 0.96, 1.43) | 1N, 1Z |
| O | Original CBA simulations (*Pure System*) | **218 ns** (0.53, 0.91, 2.03) | **241 ns** (0.33, 0.58, 0.90) | — |

[a] Frames were extracted from all OBS events identified in the original CBA simulations of the *Pure Systems*

**Table S4** Ligand-Protein and Ligand-Membrane binding free energies computed in *Pure Systems*

| Interaction | $\Delta G^{\circ}_{\text{bind}}$, kcal·mol$^{-1}$, **NECA** | $\Delta G^{\circ}_{\text{bind}}$, kcal·mol$^{-1}$, **ZMA** |
|---|---|---|
| Ligand-Protein (LP) [a] | -4.0 $\pm$ 0.2 | -5.7 $\pm$ 0.4 |
| Ligand-Membrane (LM) [b] | -1.9 $\pm$ 0.1 | -4.4 $\pm$ 0.2 |

[a] Ligand–protein binding free energy reported in this work (Main Text, Section 2.4 and Fig. 5). Computation details are provided in Methods 4.8 and Fig. S8.
[b] Ligand–membrane binding free energy reported in this work (Main Text, Section 2.4 and Fig. 5). Values represent the mean of multiple independent samples; uncertainties correspond to the standard deviation (see Fig. S9).

**Table S5** Ligand-protein binding free energies computed in mixed systems

| | System | $\Delta G^{\circ}_{\text{bind}}$, kcal·mol$^{-1}$, **NECA** | $\Delta G^{\circ}_{\text{bind}}$, kcal·mol$^{-1}$, **ZMA** |
|---|---|---|---|
| 1 | NECA 30% – ZMA 70% | -3.8 | -6.0 |
| 2 | NECA 50% – ZMA 50% | -3.7 | -5.7 |
| 3 | NECA 70% – ZMA 30% | -3.7 | -6.0 |

Binding free energies were computed as described in Section 4.8 of the Methods in the main paper.

**Table S6** Simulation box setup and molecular contents for each simulation set

| | **CGMD Simulations** | | | | **AAMD Simulations** |
|---|---|---|---|---|---|
| | **Competition 5 systems of 36** | **Unbinding 4 systems of ≥100** | **Parametr. Bonded** | **Parametr. FEP** | **Parametr. AA Ref.** |
| Box size, nm | 13 x 13 x 13 | 13 x 13 x 13 | 6 x 6 x 6 | 6 x 6 x 6 | 6 x 6 x 6 |
| Box composition [a] | WIMPL | WIMPL | WIL | SL | WIL |
| Membrane composition | POPC | POPC | — | — | — |
| Lipid molecules | 530–537 | 529–531 | — | — | — |
| Solvent molecules | 12649–13369 | 12757–12758 | 1600 | 1600 | 7000 |
| Salt ions | 198 $Na^+$, 208 $Cl^-$ | 198 $Na^+$, 208 $Cl^-$ | 20 $Na^+$, $Cl^-$ | — | 20 $Na^+$, $Cl^-$ |
| Salt conc. | 0.15 M | 0.15 M | 0.15 M | — | 0.15 M |
| Ligands in box | NECA, ZMA | NECA, ZMA | NECA/ZMA | NECA/ZMA | NECA/ZMA |
| Ligand molecules | 14 total | 2 total | 1 | 1 | 1 |
| Replicas per system | 36 | 100–120 | 1 | 1 | 1 |
| Simulation time | 30 $\mu s$ | *until unbound* | 200 ns | 200 ns | 200 ns |

[a] Acronym key: W=Water, I=Ions, M=Membrane, P=Protein, L=Ligand, S=Solvent. The label **S** is generic and can refer to any solvent. For FEP simulations (nonbonded interaction parameterization), hexadecane and octanol were used in addition to water.

**Table S7** GROMACS simulation parameters for CG and AA MD simulations

| Parameter | CGMD Simulations | AAMD Simulations |
|---|---|---|
| Cutoff scheme | Verlet | Verlet |
| Buffer tolerance, kJ/mol | 0.005 | 0.005 |
| Coulomb scheme | Reaction-Field | PME |
| Coulumb cutoff, nm | 1.1 | 1.2 |
| Epsilon r | 15 | 1 |
| VDW scheme | cutoff | cutoff |
| VDW cutoff, nm | 1.1 | 1.2 |
| Tau T, ps | 1 | 1 |
| Ref T, K | 310 | 310 |
| Tau P, ps | 12 | 5 |
| Ref P, bar | 1 | 1 |
| Minimization algorithm | steep | steep |
| Number of steps | -1 | -1 |
| Equilibration integrator | md | md |
| Delta t, ps | 0.010 | 0.002 |
| Thermostat | V-rescale | Berendsen |
| Barostat | Berendsen | Berendsen |
| Production integrator | md | md |
| Delta t, ps | 0.010/0.020 | 0.002 |
| Thermostat | V-rescale | V-rescale |
| Barostat | Parinello-Rahman | Parinello-Rahman |
| Forcefield, protein | Martini 3 | AMBER14SB |
| Forcefield, lipids | Martini 3 | LIPID17 |
| Forcefield, ligand | Martini 3 | GAFF2 |
| Forcefield, water | Martini 3 | TIP3P |

.

**Table S8** Solvent-Accessible Surface Area (SASA) data for validation of nonbonded interactions in ligand parameterization

| | **CG SASA, nm$^2$** | **AA SASA, nm$^2$** |
|---|---|---|
| Adenosine | 4.80 | 4.38 |
| NECA | 5.67 | 5.11 |
| ZMA | 6.40 | 5.95 |

.

**Table S9** *logP* values for validation of nonbonded interactions in ligand parameterization

| | **Drugbank** [a] | **CG** | **Exp.** | **ALOGPS** | **Chemaxon** |
|---|---|---|---|---|---|
| Adenosine [b] | DB00640 | -1.14 | -1.05 | -1.2 | -2.1 |
| NECA | DB03719 | -0.94 | — | -0.6 | -2.0 |
| ZMA | DB08770 | 2.67 | — | 2.83 | 2.93 |

[a] Experimental *logP* values and predicted values are taken from the Drugbank database (https://go.drugbank.com) [S1] and can be accessed using the respective DrugBank ID. [b] denosine was used as a reference scaffold due to structural similarity with NECA and ZMA, as well as the availability of experimental *logP* data.

**Table S10** Parameters used for the density function (DF) and PMF calculations in binding free energy estimation

| Parameter | Description | LP-NECA | LP-ZMA | LM-NECA | LM-ZMA |
|---|---|---|---|---|---|
| alpha | Cone aperture or apex angle, degrees | 28 | 26 | 30 | 30 |
| $R_{cutoff}$ | Boundary defining bound and unbound states, nm | 5.00 | 5.00 | 2.25 | 2.50 |
| $R_{max}$ | Distance from spherical cone apex to box edge, nm | 6.00 | 6.00 | 4.25 | 4.00 |
| d$h$ | Thickness of slices along spherical cone axis, nm | 0.25 | 0.25 | 0.50 | 0.50 |
| T | Temperature, K | 310 | 310 | 310 | 310 |

[a] For all calculations, the universal gas constant $R = 1.987 \cdot 10^{-3}$ kcal·mol$^{-1}$K$^{-1}$ and the standard concentration $C^o = 1/1.66$ nm$^{-3}$ were used.
* Abbreviation key: LP = Ligand-Protein interaction; LM = Ligand-Membrane interaction

### S3 Supplementary Discussion

#### S3.1 NECA and ZMA residence time (RT) calculations from CBA simulations

NECA and ZMA RTs were calculated directly from the original CBA simulations in pure systems and compared with the control systems reported in the main text — i.e., systems generated by removing the EXO-bound ligand from the EXO–OBS co-occupied receptor (Table 2). Specifically, RTs were determined using two complementary approaches. The first involved 100 independent unbinding trajectories initiated from conformations representative of the most populated cluster identified during the CBA, where no ligand was bound at the EXO site and the OBS-bound ligand was within 3 Å RMSD of the experimental binding pose. The second set of RTs was extracted directly from the original CBA trajectories. Both approaches yielded statistically consistent and reliable RT estimates, as confirmed by the indicators $p$-value, $m_t/m$ ratio, and $\mu/\sigma$ ratio, together with Poisson modeling (Table S3 and Fig. S4–S5). The resulting RT values were in close agreement with one another and with those obtained for the control systems, underscoring the reproducibility and robustness of our kinetic analysis protocol.

#### S3.2 EXO-OBS multiligand occupancy

The simultaneous occupation of the EXO and OBS regions represents a key feature of the multiligand binding process, playing a critical role in modulating unbinding dynamics and, consequently, ligand residence times (see Main Text, Section 2.3). From a competitive binding perspective, such conformations may correspond to a ligand poised to replace the orthosteric binder. Interestingly, our CBA simulations revealed that ligand–ligand interactions and receptor co-occupancy are not purely competitive in nature but instead reflect a nuanced interplay combining both competitive and cooperative components. Specifically, when the OBS is occupied and a second ligand resides at the EXO - effectively positioned as a potential competitor - its presence exerts a measurable influence on the dissociation behavior of the orthosteric ligand. Within this framework, we provide here a detailed structural analysis of the receptor–ligand complex under dual occupancy conditions, focusing on configurations where one ligand is bound at the OBS while another simultaneously engages the EXO site.

In detail, we performed a two-step clustering analysis on all conformations in which both the EXO and OBS sites were simultaneously occupied. The procedure involved: (1) applying a DBSCAN clustering based on the center of geometry (COG) of the EXO ligands, to identify the most frequently visited EXO subregions; and (2) conducting an RMSD-based clustering of ligand conformations within each identified subregion. From the resulting cluster families, the structure of the OBS-bound ligand exhibiting the lowest

RMSD relative to the experimental binding mode was selected and is shown in Figure 3 of the main text. In the following, we focus on the kinetically relevant conformations in which NECA or ZMA occupies the EXO site while another ZMA molecule remains bound at the OBS. As reported in the main text, the residence time of OBS-bound NECA is not significantly affected by the presence of a ligand at the EXO site; therefore, those cases are not further analyzed here. For OBS-bound ZMA, however, the cluster analysis identified three dominant EXO-binding modes for NECA (collectively representing 67% of the population) and two dominant modes for ZMA (representing 66% of the population). Representative structures of these five conformations are shown in Fig. 10.

From the figure, it is evident that the dominant EXO subregion is positioned directly above the OBS, nestled between TM6, TM7, ECL3, and ECL2. In the first EXO–NECA binding mode (Fig. S10a), the adenine ring is oriented toward ECL2, in proximity to K153$^{ECL2}$, where it can form a H-bond with K150$^{ECL2}$. The ribose moiety engages E169$^{ECL2}$, while the ethylcarboxamido group extends toward H264$^{ECL3}$. In the second mode (Fig. S10b), the ethylcarboxamido group instead interacts with ECL2, with K153$^{ECL2}$ positioned as the nearest residue, while the ribose ring remains oriented toward E169$^{ECL2}$ and T$^{6.58}$. In the third mode (Fig. S10c), the ligand shifts closer to TM6. Here, the ethylcarboxamido moiety lies near K150$^{ECL2}$, the ribose ring can form a H-bond with K153$^{ECL2}$, and the adenine group interacts with H264$^{ECL3}$. Across all three EXO–NECA binding modes, K153$^{ECL2}$ and E169$^{ECL2}$ consistently emerge as key contact residues, in agreement with previously reported features of the $A_{2A}$ vestibular site.[S2, S3] Notably, NECA's interactions with E169$^{ECL2}$ and H264$^{ECL3}$ may perturb the native ECL2–ECL3 salt bridge

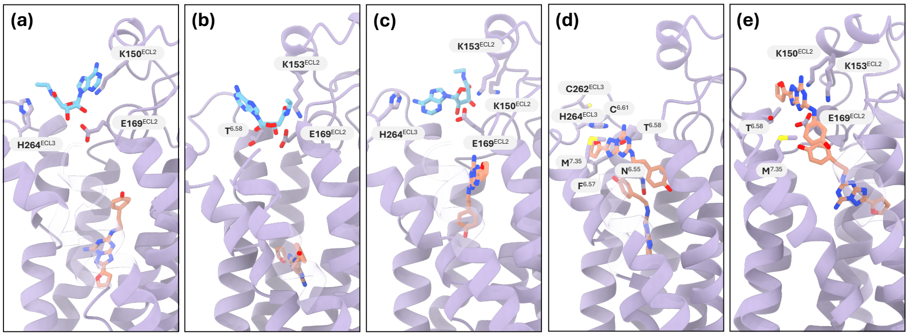


**Fig. S10 EXO binding modes of NECA and ZMA in EXO–OBS co-occupied $A_{2A}$ receptors. (a-c)** Backmapped atomistic structures of EXO-bound NECA. Shown are the centroids of the three most populated clusters identified from the ensemble of NECA–ZMA EXO/OBS co-occupancy poses, accounting for 36%, 18%, and 13% of the total population. **(d-e)** Backmapped structures of EXO-bound ZMA. Shown are the centroids of the two most populated clusters identified from the ensemble of ZMA–ZMA EXO/OBS co-occupancy poses, accounting for 56% and 10% of the total population.

formed by these residues, supporting prior evidence implicating this interaction in ligand binding mechanism and kinetics.[S2, S4–S6]

In the EXO-ZMA OBS–ZMA co-occupancy simulations, the most populated EXO–ZMA pose adopts a strikingly distinct conformation (Fig. S10e). Here, the ligand occupies a narrow crevice between TM6 and TM7 and is partially capped by ECL3. This binding geometry appears enabled by ZMA's ability to adopt a thin, elongated, highly planar architecture, allowing it to slip — slightly curved — from ECL3 (via its furan ring) down toward the upper edge of the OBS (via its phenol moiety). At the extracellular side, the furan ring interacts with the disulfide bridge $C^{6.61}$–$C262^{ECL3}$ in the highly mobile ECL3 and with $F^{6.57}$, while the adenine scaffold is stabilized by H-bonds with $T^{6.58}$ and $H264^{ECL3}$. Toward the receptor interior, the amine linker to the phenol can form H-bonds with $N^{6.55}$ and $M^{7.35}$, whereas the phenol ring itself appears relatively mobile within the cavity. The less populated second EXO–ZMA mode is located in the same vestibular subspace as the three EXO–NECA poses (Fig. S10d). Here, the furan ring can form a H-bond with $K150^{ECL2}$, along with a cation–$\pi$ interaction with $K153^{ECL2}$. With ZMA extended across the pocket, the phenol group is positioned beneath ECL3 toward TM6 and TM7, where it can interact with $M^{7.35}$. Finally, the adenine moiety is close to interact thorugh H-bonds with $E169^{ECL2}$ and $T^{6.58}$.

#### S3.3 Absolute ligand binding free energy calculation

The protocol used in this study to estimate the absolute ligand binding free energy builds upon seminal works on molecular binding affinity estimation from atomistic molecular dynamics (MD) simulations,[S7–S13] which we have further adapted for protein–ligand systems in coarse-grained MD (CG-MD) simulations.[S14] In brief, the binding free energy is derived from the one-dimensional potential of mean force, $PMF(r)$, obtained from the radial distribution function, $RDF(r)$, of a ligand around its protein. This approach characterizes the protein–ligand interaction as a function of their separation distance $r$, which serves as the reaction coordinate (see Methods 4.8 in the main text). For soluble protein–ligand systems in aqueous environments, the computed $\Delta G^o_{bind}$ values show good agreement with experimental data. However, no $\Delta G^o_{bind}$ measurements have been reported for membrane receptors.

In soluble proteins, where the ligand experiences only its interaction (or lack thereof) with the protein, a spherical reaction coordinate provides a natural and conceptually straightforward model. In contrast, for membrane proteins, the presence of the lipid bilayer introduces additional complexity. Specifically, in the $A_{2A}$ receptor system, three distinct binding equilibria coexist involving ligand (L), protein (P), and membrane (M):

$$\mathrm{L} + \mathrm{P} \xrightleftharpoons{\mathrm{K_1}} \mathrm{LP} \tag{1}$$

$$\mathrm{L + M \underset{}{\overset{K_2}{\rightleftharpoons}} LM} \quad (2)$$

$$\mathrm{LM + P \underset{}{\overset{K_3}{\rightleftharpoons}} LP + M} \quad (3)$$

Figure 5a in the main text illustrates the three-state binding model. This scheme clearly shows that the classical spherical definition of $r$ centered on the protein cannot accurately describe the protein–ligand interaction as expressed in Eq. (1). The traditional radial coordinate $r$ is indeed unsuitable for non-homogeneous environments composed of both aqueous and membrane regions. In such systems, the resulting potential of mean force (PMF) would represent the bound $LP$ state at $r = 0$ and an unbound $L$ state at $r \geq R_{cutoff}$, where the apparent $L$ density is in fact a mixture of $LM$ and $L$ states. Furthermore, the PMF would be biased by the much higher lipid concentration relative to protein in the simulation box (∼532:1 in our setup) and by the substantial volume occupied by the membrane, which is inaccessible for ligand binding. To overcome these limitations and correctly describe the solvated unbound state, we replaced the classical spherical volume with a spherical cone oriented along the receptor's principal axis and computed the ligand density function within this region. This refinement enables a clear isolation of the purely solvated $L$ state and allows a rigorous evaluation of the $L - P$binding event as defined in Eq. (1).

The resulting binding free energy thus represents exclusively the $L - P$ interaction (see Fig. 5 in the main text). Importantly, this approach inherently accounts for the membrane's influence on $L - P$ binding, as the ligand's lipid affinity contributes to its spatial density distribution in the (spherical cone) solvent. Finally, to assess the robustness of our estimates, we evaluated the effect of varying the spherical cone aperture on the computed $L - P$ binding free energy. The free-energy values showed clear convergence across cone angles, confirming the reliability of the estimates reported in the main text and in Fig. 5. The same procedure was applied to estimate the $L - M$ binding free energy, as defined in Eq. (1) and reported in Fig. 5 and Table S4.

Following the modified calculation of the distribution function, we present our thermodynamics results in Section 2.5 in the main text. The simulation set used for the measurement is only the positional data / densities from the *Pure Ligand Systems*. Nevertheless, Table S5 shows that even using data from the *Mixed Ligand Systems* lead to the same values, in line with the finding that binding events in all five original competition systems are mostl of a lone ligand environment scenario $\geq$ 97% of the time.

[S3] Hung N. Do, Sana Akhter, and Yinglong Miao. Pathways and mechanism of caffeine binding to human adenosine a2a receptor. *Frontiers in Molecular Biosciences*, 8, 4 2021.

[S4] Xueqin Pang, Mingjun Yang, and Keli Han. Antagonist binding and induced conformational dynamics of gpcr a 2a adenosine receptor. *Proteins: Structure, Function, and Bioinformatics*, 81:1399–1410, 8 2013.

[S5] Caroline Bushdid, Claire A. de March, Jérémie Topin, Matthew Do, Hiroaki Matsunami, and Jérôme Golebiowski. Mammalian class i odorant receptors exhibit a conserved vestibular-binding pocket. *Cellular and Molecular Life Sciences*, 76:995–1004, 3 2019.

[S6] Takumi Ueda, Tomoki Tsuchida, Masatoshi Kurita, Takuya Mizumura, Shunsuke Imai, Yutaro Shiraishi, Yutaka Kofuku, Shuhei Miyakawa, Kaori Fukuzawa, Koh Takeuchi, and Ichio Shimada. Structural basis of the residence time of adenosine a2a receptor ligands revealed by nmr. *Chemical Science*, 16:17948–17955, 2025.

[S7] J. E. Prue. Ion pairs and complexes: Free energies, enthalpies, and entropies. *Journal of Chemical Education*, 46:12–16, 1969.

[S8] Daniel Trzesniak, Anna Pitschna E. Kunz, and Wilfred F. Van Gunsteren. A comparison of methods to compute the potential of mean force. *ChemPhysChem*, 8:162–169, 1 2007.

[S9] E. Duboué-Dijon and J. Hénin. Building intuition for binding free energy calculations: Bound state definition, restraints, and symmetry. *Journal of Chemical Physics*, 154, 5 2021.

[S10] William L. Jorgensen. Interactions between amides in solution and the thermodynamics of weak binding. *Journal of the American Chemical Society*, 111:3770–3771, 1989.

[S11] Marie Claude Justice and Jean Claude Justice. Ionic interactions in solutions. i. the association concepts and the mcmillan-mayer theory. *Journal of Solution Chemistry*, 5:543–561, 8 1976.

[S12] E. Guàrdia, R. Rey, and J. A. Padró. Potential of mean force by constrained molecular dynamics: A sodium chloride ion-pair in water. *Chemical Physics*, 155:187–195, 8 1991.

[S13] Ilja V. Khavrutskii, Joachim Dzubiella, and J. Andrew McCammon. Computing accurate potentials of mean force in electrolyte solutions with the generalized gradient-augmented harmonic fourier beads method. *Journal of Chemical Physics*, 128:44106, 1 2008.

[S14] Paulo C.T. Souza, Sebastian Thallmair, Paolo Conflitti, Carlos Ramírez-Palacios, Riccardo Alessandri, Stefano Raniolo, Vittorio Limongelli, and Siewert J. Marrink. Protein–ligand binding with the coarse-grained martini model. *Nature Communications 2020 11:1*, 11:1–11, 7 2020.
